# Comprehensive profiling of *N*^6^-methyladnosine (m^6^A) readouts reveals novel m^6^A readers that regulate human embryonic stem cell differentiation

**DOI:** 10.1101/2025.03.09.640963

**Authors:** Zhou Huang, Rucong Liu, Zibaguli Wubulikasimu, Wanqing Zhao, Jiaqi Huang, Jiaxuan Wang, Rui Fan, Wei Kong, Qinghua Cui, Yang Li, Yuan Zhou

## Abstract

*N*^6^-methyladenosine (m^6^A) methylation has emerged as a prevalent RNA modification that extensively impacts various physiological and pathological processes via various post-transcriptional readout effects in mammals. High-throughput methylome profiling has outlined the landscape of m^6^A modification sites, but their downstream readouts require comprehensive investigation. To this end, we systematically assessed the effects of m^6^A on mRNA half-life, translation efficiency, and alternative splicing across five cell lines (A549, HEK293T, HUVEC, JURKAT, and human embryonic stem cells (hESCs)) using actinomycin D-disrupted temporal transcriptome sequencing, ribosome sequencing, and ultra-high-depth transcriptome sequencing, respectively. Our analysis, coupled with the integration of public and re-profiled m^6^A methylome data, revealed high cell type specificity in m^6^A readouts where m^6^A level alone is insufficient to predict m^6^A readouts. Machine learning models focused on the RNA binding protein (RBP) binding context of m^6^A sites demonstrated substantial predictive ability of m^6^A readouts while prioritizing putative m^6^A readers from their informative RBP features. Four novel m^6^A readers (DDX6, FUBP3, FXR2, and L1TD1) were identified and validated through m^6^A RNA pull-down assays and transcriptome-wide RBP binding site mapping. Notably, FUBP3, FXR2 and L1TD1 were found to regulate hESC differentiation without impairing self-renewal, underscoring their critical roles in stem cell biology. Together, this study bridges the gap in understanding m^6^A functional readouts and lays the groundwork for future research on m^6^A-mediated stem cell fate decisions.

## Introduction

The mammalian transcriptome is extensively regulated by a variety of chemical modifications, among which *N*^6^-methyladenosine (m^6^A) has been proven as one of the most prevalent modifications ^1–3^. The on-and-off state and quantification of m^6^A methylation are closely related to the reversible enzymatic reaction system behind. Among them, METTL3 is able to independently methylate substrate RNA *in vitro*, and *METTL3* knockdown leads to a significant reduction of transcriptome m^6^A methylation level, together highlighting METTL3 as the core component of m^6^A writer complex^4^. On the other hand, the m^6^A modification can be removed by eraser proteins such as FTO and ALKBH5 ^5^. Through sophisticated procedures executed by the m^6^A writers and erasers, a complicated epitranscriptome is formed. To date, near 500,000 human single-nucleotide m^6^A sites has been identified ^6^. Comprehensive m^6^A profiling by MeRIP-seq has revealed distinct but partly shared m^6^A profiles across different cell types ^7,8^. Nonetheless, considering the different transcriptome context among different cell types, direct and systematic measure of the regulatory effects of m^6^A is necessary. Unfortunately, there remains a lack of sizable high-throughput profiling of m^6^A’s effects, and consequently, to what extent the m^6^A regulatory effects are shared or distinct between different cell types is not clear.

The m^6^A modification is also recognized as the top RNA modification with the greatest number of known recognizing partners, i.e. m^6^A readers. This comprehensive and sophisticated reader toolkit gives rise to the diversity and complexity of the m^6^A regulatory effects, i.e. m^6^A readouts. Although with controversial about functional specificity ^9,10^, it is recognized that several key m^6^A readers play vital roles in gene expression regulations. For example, YTHDF1 would promote the translation of mRNAs with m^6^A modification, while YTHDF2 would reduce the half-life of m^6^A target transcripts ^11,12^. Nuclear YTHDC1 could recruit pre-mRNA splicing factor SRSF3 and inhibit the binding of SRSF10 to mRNAs to influence exon inclusion events in alternative splicing ^13^. IGF2BP family proteins constitute a new clade of m^6^A readers that inhibit mRNA degradation and enhance translational efficiency ^14^. Nonetheless, disruption of any one of these m^6^A readers cannot completely abolish even single type of regulatory effects of m^6^A ^11–14^, suggesting the existence of undiscovered m^6^A readers. Indeed, by leveraging mass spectrometry, dozens of proteins have shown favorable presence in the m^6^A-associated proteome, although the direct m^6^A recognition ability of these putative m^6^A readers need further validations ^15^. Until recent, novel m^6^A readers have been unveiled. For example, ELAVL1 (HuR) preferentially binds elements in the 3’-UTR that are enriched for AU or U, thereby regulating mRNA stability ^16^. TARDBP (TDP43) is identified as a m^6^A-binding protein that regulate neurodegeneration ^17^.

As the key epitranscriptomic factor, m^6^A also exert substantial impacts on key properties of pluripotent stem cells. In fact, early demonstrations of m^6^A regulatory function on embryonic stem cells (ESCs) constitute the pioneer cases supporting the functional importance of m^6^A modification. In 2014, Wang et al. reported that *Mettl3* inhibition decreases the expression of pluripotency-associated genes (e.g., *Nanog* and *Sox2*) but up-regulates the expression of developmental markers (e.g., *Sox17*), and ultimately promotes mouse ESCs (mESCs) differentiation ^29^. In induced pluripotent stem cells (iPSCs), Chen et al. also showed that increased m^6^A formation promotes cell reprogramming to pluripotency ^18^. However, there are also inconsistent results, for example, in 2014, Batista et al. found that depletion of mouse and human *Mettl3* promoted self-renewal of mESCs and human ESCs (hESCs) by prolonging *Nanog* expression upon differentiation. Subsequently, Geula et al. showed that deletion of m^6^A could lead to either defected or accelerated differentiation depending on the pluripotent state of the mESCs, i.e. naive or primed ^19^. These results imply complexity in the functional readout of the m^6^A tags in ESCs. Besides, various m^6^A readers also play essential roles in regulating ESC pluripotency. Knockout of *YTHDF* genes inhibits mESCs exit from pluripotency as evidenced by poor differentiation during teratoma and embryoid body formation ^20^. Deletion of YTHDC1 initiates cellular reprogramming to a 2-cell like state^21^. These studies emphasize the importance of the m^6^A modification in regulating the self-renewal and differentiation process of ESCs. However, which and how m^6^A readers exert the specific regulatory effects of hESC differentiation has not been sufficiently investigated.

Together, m^6^A has emerged as the key gene expression regulatory machinery but its regulatory effects and mechanisms have not been fully elucidated. Whether m^6^A showed similar or distinct readouts across different cell types? Are there new readers behind the regulatory effects of m^6^A. And if so, how could these m^6^A readers influence basic stem cell functions including self-renewal and differentiations? To this end, we comprehensively profiled the three main layers of m^6^A readouts (mRNA half-life, translational efficiency and alternative splicing) across five common cell types of different origin (including cancerous A549 and JURKAT, and non-cancerous hESC, HEK293T and human umbilical vein endothelial cells (HUVECs)). Besides, quantitative profiling of m^6^A methylation by GLORI, if applicable, was also performed. To exploit these resources, machine learning modeling of the m^6^A readouts was conducted, where putative m^6^A readers were suggested based on the interpretation of model features. Finally, we validated a few of these putative m^6^A readers and unveiled their specific impacts on hESC differentiation.

## Methods

### Cell culture

HEK293T and A549 cell lines were cultured in DMEM medium (Gibco, catalog#C11995500BT) supplemented with 10% fetal bovine serum (FBS, Gemini, catalog#900-108) and antibiotics. JURKAT cell line was maintained in RPMI 1640 medium (Gibco, catalog#C11875500BT) with 10% FBS (Gemini, catalog#900-108) and antibiotics. hESCs were cultivated in serum-free, defined mTeSR1 medium (STEMCELL, catalog#85850) and grown on plates coated with growth-factor-reduced Matrigel. hESCs were passaged with non-enzymatic dissociation by 0.5 mM EDTA (Beyotime, catalog#C0196-100ml). HUVECs were isolated as described ^22^. The clean vein of the umbilical cord was first lavaged with 0.25% trypsin at 37°C for 15 min. HUVECs were collected and suspended in Endothelial Cell Medium (ECM, ScienCell, catalog#1001). Isolated HUVECs were cultured between passage 4–6 in ECM for experiment. Cells were grown at 37°C with 5% CO2.

### *METTL3* knockdown, transfection and flow cytometry

The doxycycline-controlled Tet-Off/On gene expression systems were used for regulating the activity of METTL3 in cell lines. The sequence targeting *METTL3* were synthesized and inserted into the lentivirus vector GV307 (TetIIP-TurboRFP-MCS-Ubi-TetR-IRES-Puromycin) and the lentiviruses were purchased from GeneChem (Shanghai, China). Then according to the characteristics of different cell lines, lentiviruses transfection was performed using the HiTransG A/P developed by GeneChem. Cells transfected with shRNA lentivirus or negative control lentivirus were cultured for 48 h. The target sequence of *METTL3* was CAAGGAACAATCCATTGTT and the negative control sequence of insertion was TCTCGCTTGGGCGAGAGTAAG. Other information of the oligo synthesis was included in **Supplementary Table S1**. After the doxycycline induction for 48 h, the transfected cells expressed red fluorescence (RFP) for roughly estimating the overall transfection efficiency. Cell samples were resuspended in MACS buffer for fluorescence-activated cell sorting (FACS, BD FACSAria™ III) to obtain high RFP expression transfected cells.

### RNA extraction

Total RNA was isolated from cell samples using TRIzol (Invitrogen, catalog#15596018) following the manufacturer’s instruction. Briefly, cells were lysed with TRIzol reagent at room temperature until the cell deposits disappear and chloroform was added. After centrifuge for 15 min at 12,000×g at 4°C, the aqueous phase was transferred to the new EP tube, mixed with isopropanol and stand at room temperature for 10∼15 min. Then centrifuge for 10 min at 12,000×g at 4°C, the supernatant was discarded and the RNA precipitate was washed with 75% ethanol. After centrifuge for 5 min at 7,500×g at 4°C and dry, RNA was dissolved with RNase-free water and its concentration was measured by Thermo Scientific NanoDrop.

### Real-time qPCR for mRNA quantification

Total RNA was extracted from transected cells scribed in ‘RNA-extraction’, then reverse transcribed using PrimeScript RT Master Mix (TaKaRa, catalog#RR036B). All primers used for RT-qPCR were listed in **Supplementary Table S2**. All RT-qPCRs were run on Bio-Rad CFX384 real-time PCR detection system using PowerUp SYBR Green Master Mix (ThermoFisher Scientific, catalog#A25742). GAPDH was used as internal control to normalize the data across different samples.

### Western blot

Cell samples were collected, washed with ice-cold PBS and fully lysed with RIPA lysis buffer containing protease inhibitors (Beyotime, catalog#P0013C) on ice. After centrifuge for 30 min at 12,000×g at 4°C, supernatant was collected and protein amounts were quantified using the BCA Protein Assay Kit (Beyotime, catalog#P0012S). After boiling for 10 min, equal amounts of protein were loaded and separated by 10% SDS-PAGE. After electrophoresis, they were transferred to PVDF membranes (Merck-Millipore, catalog#ISEQ00010). Membranes were blocked with 5% non-fat milk (Beyotime, catalog#P0216-1500g) for 1 h and then incubated at 4°C overnight with the primary antibodies. After incubating 1 h with secondary antibodies at room temperature, images were obtained using the Gel Doc EZ Imager (Bio-Rad). GAPDH was used as the loading control. All antibodies used for western blot were listed in **Supplementary Table S3**.

### High-throughput profiling of m^6^A readouts

Unless otherwise stated, the m^6^A readouts were measured by comparing shControl (normal m^6^A level) versus shMETTL3 (disrupted m^6^A level) to intuitively depict the regulatory readouts in presence of m^6^A. The sequencing service related to m^6^A readout estimations was provided by Novogene Biotech (Tianjin, China). The experimental and computational pipelines of each readout category is described as follows:

#### a. mRNA half-life readout

Cells transfected with *METTL3* shRNA or negative control shRNA were re-seeded into 6 well plates. After the doxycycline induction for 48 h, actinomycin D was added to 2 μg/mL at 0 h, 1 h, 3 h and 6 h before total RNA extraction. Before construction of the library with mRNA sample, ERCC RNA spike-in control mix (Invitrogen, catalog# 4456740) was added to each sample following the manufacturer’s instruction. NEBNext Ultra RNA Library Prep Kit for Illumina (NEB #E7770) and NEBNext Ultra Directional RNA Library Prep Kit for Illumina (NEB #E7760) were used with 400 ng of total RNA for the construction of sequencing libraries. ERCC spike-in were added to total RNA before library construction, following the protocol recommended by ERCC spike-in kit manufacturer. RNA libraries were prepared for sequencing using standard Illumina protocols. Two biological replicates were prepared for each time point. For each sample collected at 1 h, 3 h and 6 h, a 15G amount of paired-end sequencing reads was obtained. For 0 h sample, we required 60G amount of sequences to ensure ultra-deep RNA sequencing that is necessary for systematic alterative splicing assay. This differences in data size were properly controlled by the aforementioned ERCC spike-ins. More specifically, quality control, adapter cutting and low-quality sequence removal were performed by FastQC, fastp and FASTX toolkits. The clean reads were mapped to human genome (hg38) by HISAT2 software (v2.2.1) and the expression values were calculated by featureCounts command of RSubread package with suggested parameter of the software. The absolute quantification of each sample was performed by using ERCC RNA spike-ins as the reference, following Wang et al. ^11^. The mRNA half-life was also estimated following the method modified from ^11^. The degradation rate of RNA *k* was estimated by:

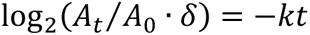

where *t* is transcription inhibition time (h), 𝐴_𝑡_ and 𝐴_0_ represent mRNA quantity at time *t* and time 0, 𝛿 is a regulating factor in case the RNA:ERCC ratio is prominently deviated from the expectation. *k* values of 3 h and 6 h were calculated: time 3 h versus time 0 h, and time 6 h versus time 0 h. A 1.5-fold change of half-life threshold was applied for subsequent analysis.

#### b. translational efficiency readout

The translational efficiency was measured by ribosome profiling sequencing (Ribo-seq). The reported protocols ^23^ was adapted with the following modifications: Before cells collection, cycloheximide (CHX, HARVEYBIO, catalog#C21865-100mg) was added to the culture media at 100 μg/mL for 5 min. Eighty million cells from each group were collected, rinsed in ice-cold PBS with 100 μg/mL CHX. After centrifuge for 5 min at 500×g at 4°C, cells were resuspended in cell freezing medium with 100 μg/mL CHX for dry ice transportation. Two replicates were prepared and the reads were processed and analyzed with the similar pipeline to the above part a. To comply with the Ribo-seq convention, the R1 reads were used for subsequent analysis. The Xtail software (v1.1.5) was utilized to measure the changes of translational efficiency with default parameter ^24^. To ensure sufficient number of genes for subsequent analysis, a relaxed thresholds of 1.5-fold change and P < 0.05 were applied. We also considered public Ribo-seq data from GEO database for the machine learning modeling described later, including those from HeLa (GSE117299), HepG2 (GSE121952), Huh7 (GSE155447) and MOLM13 (GSE98623). These data were analyzed with the same pipeline.

#### c. alternative splicing readout

We exploited the ultra-deep sequencing of 0 h samples as described in the above part a to measure alternative splicing events. Two replicates were prepared and the reads were processed and analyzed with the similar pipeline to the above part a. The alternative splicing events and exon inclusion ratio (Psi) were calculated by rMATS-turbo (v4.1.0, default parameter) ^25^. A change of exon inclusion ratio (delta-Psi) no less than 0.1, together with FDR < 0.05, were selected as the significant alternative splicing event.

#### d. Functional analysis

The functional enrichment of Gene Ontology (GO), Kyoto Encyclopedia of Genes and Genomes (KEGG), WikiPathways and MSigDB functional gene sets were performed by ClusterProfiler with default parameter and cutoffs ^26^. Gene dependency score was obtained from DepMap database ^27^.

### High-throughput profiling of m^6^A methylation sites

#### a. GLORI sequencing

Recently, Liu et al. developed GLORI, an antibody-independent sequencing technique for single-base quantitative detection of m^6^A methylation ^28^. The main idea of GLORI is to find a catalytic system of glyoxal and nitrite through chemical reaction combination screening, which efficiently deaminate unmethylated adenosines to inosines (A-to-I), but not methylated ones. We used this state-of-the-arts technique for the quantitative profiling of m^6^A methylation in A549, hESC and JURKAT. The GLORI-identified m^6^A sites in HEK293T was directly obtained from the original study ^28^. HUVEC was not included because of the required large amount of input cells by GLORI sequencing service. GLORI sequencing service was provided by Cloud-Seq Biotech (Shanghai, China). Total RNA from the samples was extracted using TRIzol, and RNA was purified according to the MEGA clear transcription purification kit (Thremo fisher, catalog#AM1908). RNA was incubated with fragmentation buffer (NEB, catalog#E6150S) for 4 min at 94°C and glyoxal solution and DMSO were added, and the mixture was incubated for 30 min at 50°C in a preheated thermal cycler. minutes, and at the end of the incubation, the tubes were brought to room temperature, and then saturated H3BO3 (Sigma-Aldrich, catalog#B0394) solution was added. The incubation was continued at 50 °C for 30 min, and was then added to the pre-configured deamination buffer (750 mM NaNO2 (Sigma-Aldrich, 31443), 40 mM MES (pH 6.0), and 10 μL of glyoxal solution for a total of 8 h of incubation. Finally, RNA was purified by ethanol precipitation, and sequencing libraries were constructed by GenSeq Low Input RNA Library Prep Kit (GenSeq, Inc.) following the process of the instructions provided by the manufacturer. The constructed sequencing libraries were subjected to quality control and quantification by BioAnalyzer 2100 system (Agilent Technologies, USA), followed by 150 bp paired-end sequencing.

#### b. GLORI data analysis

Quality control of raw reads was first performed using Q30. Then cutadapt software (v1.9.3) was used for read de-multiplex and removal of low-quality reads. PCR duplicates were removed based on Unique Molecular Identifiers (UMI) tags. Clean reads were aligned to the reference genome (hg38) and BAM files were obtained using STAR (v2.7) and bowtie2 (v2.2.4) software. To ensure acceptable m^6^A site detection rate, we performed two rounds of sequencing, and the aligned BAM files from these two rounds were combined by samtools (v1.12). Finally, the m^6^A sites were called and quantified by GLORI-tools ^28^ with parameter “--combine --rvs_fa -c 1 -C 0 -r 0.1 -p 0.05 -adp 0.05”. Finally, only m^6^A sites that showed modification rate > 0.1 for both replicates were retained ^28^.

#### c. Public MeRIP-seq methylation profiles

Currently, GLORI methylation profiles are not available for most cell lines. To obtain m^6^A sites for other cell types, we exploited widely used MeRIP-seq-based m^6^A methylation profiles ^1,2^. To ensure comparable results, raw sequences of MeRIP-seq studies were obtained from GEO (as specified in **Supplementary Table S4**). The adapter trimming and quality control was performed by fastp, and the reads were aligned to human genome (hg38) by HISAT2. Finally, exomePeak2 was used to call m^6^A peaks ^6,29^. To obtain single-nucleotide m6A sites, we compared m6A peaks with known single nucleotide m^6^A sites from m6A-Atlas V2.0 database ^6^ and GLORI technique by using bedtools (v2.31.0). Sites following the consensus m^6^A motif DRACH were retained.

### Machine learning modeling of readouts

#### a. Obtaining RNA binding protein (RBP) binding site annotations

Our m^6^A readout prediction models were heavily relied on transcriptome-wide annotation of RBP binding sites. CLIP-seq and its derivates constitute the major source of such annotation. We obtained CLIP-seq (and derivates)-based RBP binding sites from POSTAR2 database ^30^. We did not consider POSTAR3 because there was no RBP binding site score in the downloadable data of POSTAR3. Moreover, to make our annotations up-to-date, we obtained and re-analyzed raw CLIP-seq data from GEO, and adopted a pipeline similar to that of POSTAR database to process these data. Briefly, raw reads were filtered and trimmed by fastp and cutadpat software, and aligned to human genome (hg38) by using bowtie (v1.3.0, for short reads) or STAR (for longer reads). Samtools was used to process the aligned reads and the RBP binding regions were called by Piranha (v1.2.1) with recommended parameters ^31^. We also obtained eCLIP-based annotation from ENCODE projects ^32^. We noted that the default merging of biological replicates in ENCODE would result in very limited binding sites for some RBPs. To this end, we also used bedtools to obtain the union of RBP binding regions derived from both replicates to enhance the sensitivity and transcriptome coverage. Finally, RBP site annotations from non-CLIP techniques were directly obtained from the corresponding GEO accession. The summary of RBP binding site profiles were available in **Supplementary Table S5**.

#### b. Feature encoding and model training

For predicting half-life and translational efficiency readouts, eleven set of features were considered: 1) The score of RBP binding site with the nearest distance on the genome; 2) The score of RBP binding site with the nearest distance on the transcripts; 3) The minimum RBP-to-m^6^A-site distance among different m^6^A sites of the same gene on the genome; 4) The minimum RBP-to-m^6^A-site distance on the transcripts; 5) The maximum RBP-to-m^6^A-site distance on the genome; 6) The maximum RBP-to-m^6^A-site distance on the transcripts; 7) The site to gene topology (e.g. minimum distance to start codon); 8) Tissue expression level, as measured by GTEx project ^33^; 9) Gene dependency score, as record by DepMap database ^27^; 10) Nearest distance to a 6-mer on transcript; 11) Maximum counts of a specific k-mer in ±100 nt range flanking known m^6^A sites. As for the prediction of exon inclusion ratio alterations, we used the annotation of the closest m^6^A site to the exon boundary to encode features (if any within 10kb of exon boundary). Since only one representative m^6^A site was considered for each exon, no minimum/maximum operation of RBP features among multiple m6A sites on the same gene was required. Besides, as an exon-level prediction, gene features like GTEx tissue expression and DepMap gene dependency score were also not included.

#### c. Machine learning model

We exploited XGBoost ^34^ to model the readouts for each readout and each cell type. XGBoost has been successfully implemented in both bioinformatic prediction and clinical application ^35,36^. Ten-fold cross-validation to train and test the model, and the overall performance was measure by receiver operating characteristic (ROC) curve and area under ROC curve (AUC). A heustic search of best combination of feature set was performed to measure the AUC gain of the feature set. And for RBP features, the feature importance score derived from the XGBoost models.

### RNA pull-down

The RNA oligonucleotides [ss-A: 5’-CGUCUCGGACUCGGACUGCU-3’; ss-m^6^A: 5’-CGUCUCGG(m^6^A)CUCGG(m^6^A)CUGCU-3’] with the same sequences of ss-A and ss-m^6^A were synthesized by Rui Biotech (Beijing, China). RNA pull-down assay was conducted by the Pierce™ Magnetic RNA-Protein Pull-Down Kit (Thermo Fisher Scientific, catalog#20164). The included Thermo Scientific Pierce RNA 3’ Desthiobiotinylation Kit (catalog#20163) was used to label the target RNA. Next, biotin-labeled RNA oligonucleotides containing adenosine or m^6^A (50 pmol) were immobilized onto 50 μL streptavidin magnetic beads in binding buffer at room temperature for 15-30 min. RNA-conjugated streptavidin beads were then incubated with 200 μg total proteins from target cells in binding buffer in a final volume of 100 μL at 4°C for 30-60 min. RNA-protein complex-containing beads were washed three times with wash buffer. After extensive washing, RNA-binding protein complexes were boiled and eluted in 1 × SDS loading buffer to separate on 10% SDS-PAGE gels for western blot analysis.

### RBP binding site profiling by HyperTRIBE

#### a. HyperTRIBE plasmids construction and transfection

HyperTRIBE is a promising high-throughput sequencing technique for identifying potential binding site of an RBP. Leveraging plasmid fusing target RBP with hyperactive ADAR, the RNA editing signals close to the RBP binding site can be detected at transcriptome level through conventional RNA sequencing ^37^. Here, the hyperactive ADAR containing TRIBE plasmids were constructed by Rui Biotech (Beijing, China). The control plasmid, mammalian expression plasmid, containing mCherry ADAR control and p2A GFP reporter can be obtained from Addgene (Plasmid #154786). As for RBP-TRIBE plasmid, using standard restriction enzyme-based cloning to clone known and putative m^6^A reader RBPs (YTHDF1, DDX6, FUBP3, FXR2 and L1TD1) into TRIBE plasmid. Sanger sequencing was used to confirm proper insertion and then control and RBP-ADAR plasmids were infected into hESCs by transfection reagent jetOPTIMUS (Polyplus, catalog#101000051). After transient transfection into target cells for 24 h, the cells containing RBP-ADAR fusion or ADAR alone proteins were sorted by GFP and mCherry fluorescence and using TRIzol reagent to extract the total mRNA from the sorted cells.

#### b. TRIBE sequencing and analysis

RNA-sequencing for cells transfected with HyperTRIBE plasmid were conducted by Novogene Biotech (Tianjin, China). Messenger RNA was purified from total RNA using poly-T oligo-attached magnetic beads. After fragmentation, the first strand cDNA was synthesized using random hexamer primers followed by the second strand cDNA synthesis. After library quality control, quantified libraries were pooled and sequenced on Illumina NovaSeq with paired-end 150-bp reads, providing sequencing data at 40G per library. According to Biswas et al.’s protocol ^38^, firstly, cutadapt was used to remove the adaptor and low-quality reads. Clean reads were then aligned to the reference genome (hg38) by using STAR software. Picard (v3.3.0) was used to remove duplicates and sort the output BAM files. To identify editing sites, we used Bullseye, a set of Perl scripts which was originally developed by Flamand et al. for analysis of editing events induced by transfected plasmids ^9^. BAM files were parsed to generate coverage matrices at each position in the genome, excluding duplicate and multi-mapped reads. Editing sites for HyperTRIBE in individual samples were then called by comparing A2I mutation rates at all positions within annotated exons (hg38) with samples expressing ADAR alone. Sites edited at rates between 5% and 99%, edited to higher levels than control cells (1.5 × for A2I), with at least two mutations, and that did not overlap annotated SNP were kept for further analysis. The final list of sites was identified as those found in three replicates and with an average editing rate of at least 5% in all samples where sites were called. To identify the binding motif of the editing sites, we broadened each site by 500 bp in each direction and then imported into HOMER software ^39^ for *de novo* motif identification. It is worth mentioning that because we are interested in identification of m^6^A-binding proteins, we further focused on the proteins that have motifs containing ‘RRACH’, which represents the most common m6A consensus sequence. To investigate the binding preference of the RBPs near m6A sites, BAM alignments were further processed using deepTools ^40^. Heatmaps and profiles were generated using the plotHeatmap and plotProfile script from the deepTools suite.

### CRISPR-mediated *FUBP3*, *FXR2* and *L1TD1* knock out (KO)

The CRISPR-Cas9 knock-out vector, containing cassettes for GFP, Cas9 endonuclease, and the double guide RNA (gRNA) moiety designed to target FUBP3, FXR2 and L1TD1, was custom synthesized by Guangzhou Regenverse Therapeutics Co., Ltd. (Guangzhou, China). After plasmid transfection for 48 h, GFP+ hESCs were sorted using flow cytometry, and plated at low density for single-colony isolation. Selected single colonies were tested by western blot for loss of protein. Mutations were validated by sequencing products of PCR amplification of the regions flanking the targeting sites.

### Cell proliferation assays

#### a. Cell counting kit 8 (CCK8) assay

Human embryonic stem cells (hESCs) were digested into single cell suspension using Accutase (Merck-Millipore, catalog#SCR005) and the number of cells were counted by automated cell counter (RWD, catalog#C100). After counting, the cells were seeded in Matrigel-coated 96-well with a cell density of 2000 cells per well. After appropriate cultivate time, 10% CCK8 (Dojindo, catalog#CK04) was added into the culture medium and incubate cells for 2 h, then optical density value at 450 nm was detected using a microplate reader. Cells were detected in every 24 h.

#### b. Alkaline phosphatase staining assay

For ALP (alkaline phosphatase) staining, hESCs were digested into single cell suspension using Accutase and the number of cells were counted by automated cell counter. After counting, the cells were seeded in Matirgel-coated 12-well with the cell density of 1.2 × 10^5^ cells per well and incubated in mTeSR1 medium for 3 days. Then the cells were fixed with 4% paraformaldehyde for 3–5 min followed by washing with PBS. Next, the samples were added with BCIP/NBT staining solution (Beyotime, catalog#C3206) for 15 min in dark at room temperature. The usage of regents was followed with the manufacturer’s recommendations. The cell images were taken after washing the cells with PBS.

### hESCs differentiation assays

#### a. Endodermal differentiation

To initiate endodermal cell differentiation, hESCs were cultured on Matrigel-coated 6-well plate in RPMI1640 basal medium supplemented with B27 (Gibco, catalog#12587-010) and 100 ng/ml Activin A (Peprotech, catalog#AF-120-14E-10UG). After cell differentiation inducing for 3 days, endoderm markers were detected by flow cytometry and RT-qPCR analysis.

#### b. Hematopoietic mesodermal differentiation

For Hematopoietic mesodermal differentiation, hESCs were cultured to 60% density on Matrigel-coated plates. Using Accutase to digest cells for 3-5 min, the cells were resuspended by blowing with mTeSR1 and seeded into ultra-low attachment 6-well plates with the cell density of 2× 10^5^ cells per well. Cells were cultured in a hypoxic incubator (5% CO2, 5% O2) for 24 h, until they formed to embryoid bodies (EB). To induce mesodermal differentiation, STEMdiff APELTM2 (STEMCELL, catalog#05270) was used as the basal medium, and the differentiation medium was configured by adding corresponding cell factors according to the Kaufman hematopoietic differentiation protocol ^41^. After cell differentiation inducing for 6 days, mesodermal markers were detected by flow cytometry and RT-qPCR analysis.

#### c. Neural ectodermal differentiation

Monolayer culture protocol was used to initiate neural ectodermal differentiation. hESCs were plated onto Matrigel-coated 6-well plates with the cell density of 2 × 10^5^ cells/cm^2^, and then cultured in STEMdiff Neural Induction Medium (STEMCELL, catalog#08581) added with SMADi. After cell differentiation inducing for 3 days, neural ectodermal markers were detected by flow cytometry and RT-qPCR analysis.

### Statistical analysis

Unless otherwise stated, bar charts represent the mean ± standard deviation of the mean (SD), and the results for the two groups were compared using a two-tailed unpaired Student’s *t*-test. A *P*-value smaller than 0.05 was considered significant unless stated differently, and the exact degree of significance as indicated by asterisks is stated in the legends. Statistical significance was presented as \**P* < 0.05, \*\**P* < 0.01, \*\*\**P* < 0.001 and \*\*\*\**P* < 0.0001; ns indicates *P* > 0.05.

## Results

### Systematic profiling of m^6^A methylation readouts across different cell types

The overall design of this study is illustrated in **Figure 1A**. Briefly, we perturbed m^6^A in five representative cell lines through knockdown of core m^6^A writer *METLL3*. The efficiency of the knockdown was verified by qRT-PCR and Western blot (**Supplementary Figure S1)**. By procedure, we could measure the mRNA half-life, translation efficiency, and alternative splicing readouts of m^6^A by comparing the post-transcriptional gene expression dynamics of m^6^A-normal cells (i.e. shControl cells) versus m^6^A-disrpted cells (i.e. shMETTL3 cells). Actinomycin D-treated temporal RNA-seq, Ribo-seq and ultra-deep RNA-seq were adopted to measure the three main categories of m^6^A readouts, respectively. We introduced a cost-effective experiment design by performing ultra-deep RNA-seq on the untreated (0h) cells, which could also serve as the start point for temporal RNA-seq as well as the input sample for Ribo-seq, with the discrepancy in sequencing library size controlled by ERCC spike-ins (see Methods).

**Figure 1.**
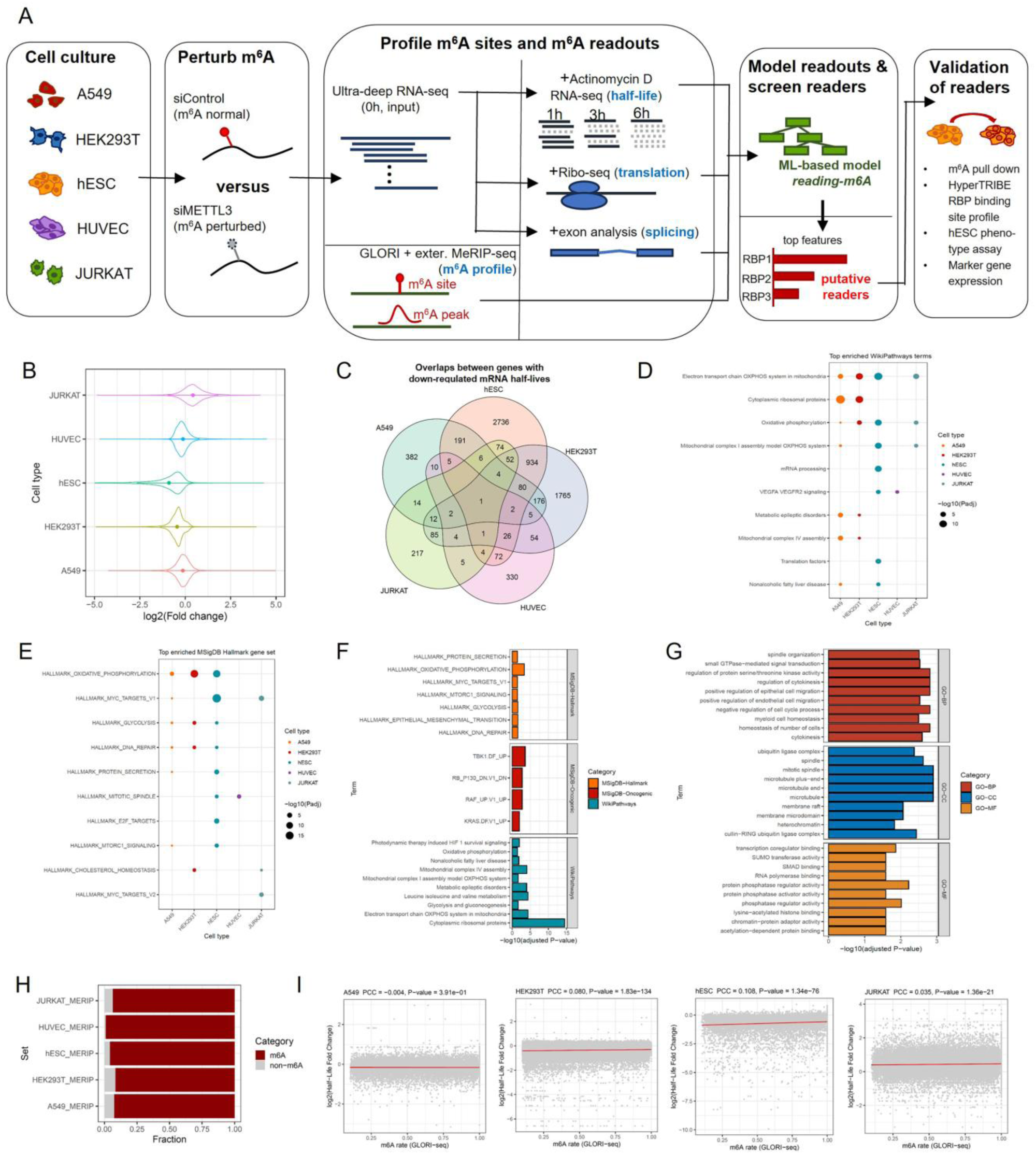
Overview of the experimental design for m^6^A readout profiling and the half-life readout profiling results. (A) Overview of the experimental design for m^6^A readout profiling. (B) Overview of m^6^A-mediated half-life changes. The comparison was performed between m^6^A-normal cells (shControl) versus m^6^A-disrupted cells (shMETTL3) as the background. (C) Venn diagram of half-life down-regulated genes in different cell types. (D) Top 10 enriched pathways (WikiPathways) of half-life down-regulated genes among different cell types sorted by total -log10(FDR). (E) Top 10 enriched MSigDB Hallmark gene sets of half-life down-regulated genes among different cell types. (E) Top enriched pathways and gene sets for half-life down-regulated genes in A549 cells. (G) Top enriched GO functional terms for half-life down-regulated genes in HUVEC cells. (H) Percentage of half-life down-regulated genes with m^6^A methylation peaks identified by MeRIP-seq assay on the corresponding cell types. (I) Correlation between fold change in half-life and m^6^A methylation levels assessed by GLORI.

The mRNA half-life was firstly measured by fitting the gene expression to the mRNA decay curve along the time points. For all cell types, a decrease of absolute mRNA quantification along the time points were observed, demonstrating substantial mRNA degradation with the transcription process inhibited by Actinomycin D. Nonetheless, the decay rate of the mRNA half-life could be diverged between different m^6^A states, where for most of the cell types, an overall down-regulation of mRNA half-lives in m^6^A normal cells (shControl cells) could be observed (**Figure 1B**), which is consistent with previous literature ^11,42^. Nonetheless, the overlap of half-life down-regulated genes is limited between different cell types, indicating a high cell-type specificity of the half-life readout of m^6^A (**Figure 1C**). It is possible that the limited overlap between down-regulated genes is simply caused by the threshold in measuring the readout. Thus, we performed enrichment for different functional gene sets and pathways to check if different cell types could show consistent changes at the pathway level. Indeed, gene sets representing fundamental biological functions such as mitochondrial oxidative phosphorylation, glycolysis and DNA repairs were shared by multiple cell types, indicating a consensus regulation by m^6^A in these basic cellular functions (**Figure 1D-E**). On the other hand, some pathways were only enriched in particular cell type, and these pathways are likely to be associated the functions of specific cell type. For example, multiple oncogenic pathways were enriched in lung cancer cell line A549, whereas cytoskeletal and endothelial cell migration functions were enriched in blood vessel-derived HUVEC cells (**Figure 1F-G**).

By querying to the public MeRIP-seq methylation profiles from the corresponding cell type, we confirmed that most (>90%) of the half-life down-regulated genes were the direct target genes of m^6^A methylation (**Figure 1H**). To precisely describe the relationship between m^6^A modification levels and half-life changes, we evaluated the m^6^A methylation levels of target genes in different cells using GLORI technique ^28^. The quantitative m^6^A map of four out of the five cell types were included, where A549, hESC and JURKAT data were newly measured in this study. HUVEC was not included in GLORI analysis because of its proliferation capacity is insufficient to accumulate required amount of input cells. The m^6^A methylation sites identified by GLORI were all enriched near the stop codon (**Supplementary Figure S2A**), which is a typical topology feature of m^6^A sites on coding genes ^2^. Our GLORI experiments could capture m^6^A sites with medium-to-high modification ratios, although low rate sites were relatively underrepresented because of the reduced sequencing depth due to the cost concerns (**Supplementary Figure S2B**). Besides, the m^6^A sites and genes were partially shared between different cell types, and noticeable overlaps between m^6^A target genes identified by GLORI and those from MeRIP-seq could be observed, confirming that the main m^6^A target genes were captured (**Supplementary Figure S2C-E**). Finally, >80% of the half-life down-regulated genes harbored a GLORI-derived m^6^A site (**Supplementary Figure S2F**), enabling a direct correlation analysis between the absolute quantification of m^6^A sites and the mRNA half-life fold-change of their target genes. Significant correlations between m^6^A modification rates and the half-life changes in most cells were observed, suggesting the half-life regulation would partly rely on the methylation level of the target genes. Nonetheless, the correlations were generally weak (**Figure 1I**), suggesting that other factors, such as the genomic context of m^6^A sites, may also have significant contributions.

Next, we profiled the translational efficiency readout of m^6^A. In terms of translational efficiency, different cell types showed more balanced changes, with roughly comparable numbers of up- and down-regulated genes (**Figure 2A**). The overlap of genes showing translational efficiency changes was also very limited between different cell types, for both up- and down-regulated genes (**Figure 2B-C**). Some of the genes showed both half-life and translational efficiency changes (**Figure 2D and Supplementary Figure S3A-C**). In A549 cell lines, genes with half-life or translational efficiency changes had higher importance (DepMap dependency ^27^) scores than background, and those with both types of changes ranked the highest; in JURKAT, higher importance scores of m^6^A-regulated genes were also observed (**Figure 2E**). Similar to the observation in half-life readout analysis, some fundamental pathways are also the favored target of the translational efficiency regulations of m^6^A (**Supplementary Figure S3D-I).** Finally, more than 90% of translational efficiency-change genes were also the m^6^A methylation target genes, as suggested by the MeRIP-seq methylation profiles (**Figure 2F**). When considering GLORI-based methylation profiles, approximately 80% of genes showing translational efficiency changes could be targeted by m^6^A (**Supplementary Figure S2G**), but GLORI-estimated methylation levels were only weakly correlated or non-significantly correlated with the fold change in translational efficiency (**Figure 2G**). We also noted that genes with up- or down-regulated translational efficiency did not always tend to have a m^6^A site near the start codon (**Figure 2H**). These results indicate that neither m^6^A level nor m^6^A topology is sufficient to fully explain the translational efficiency readout of m^6^A.

**Figure 2.**
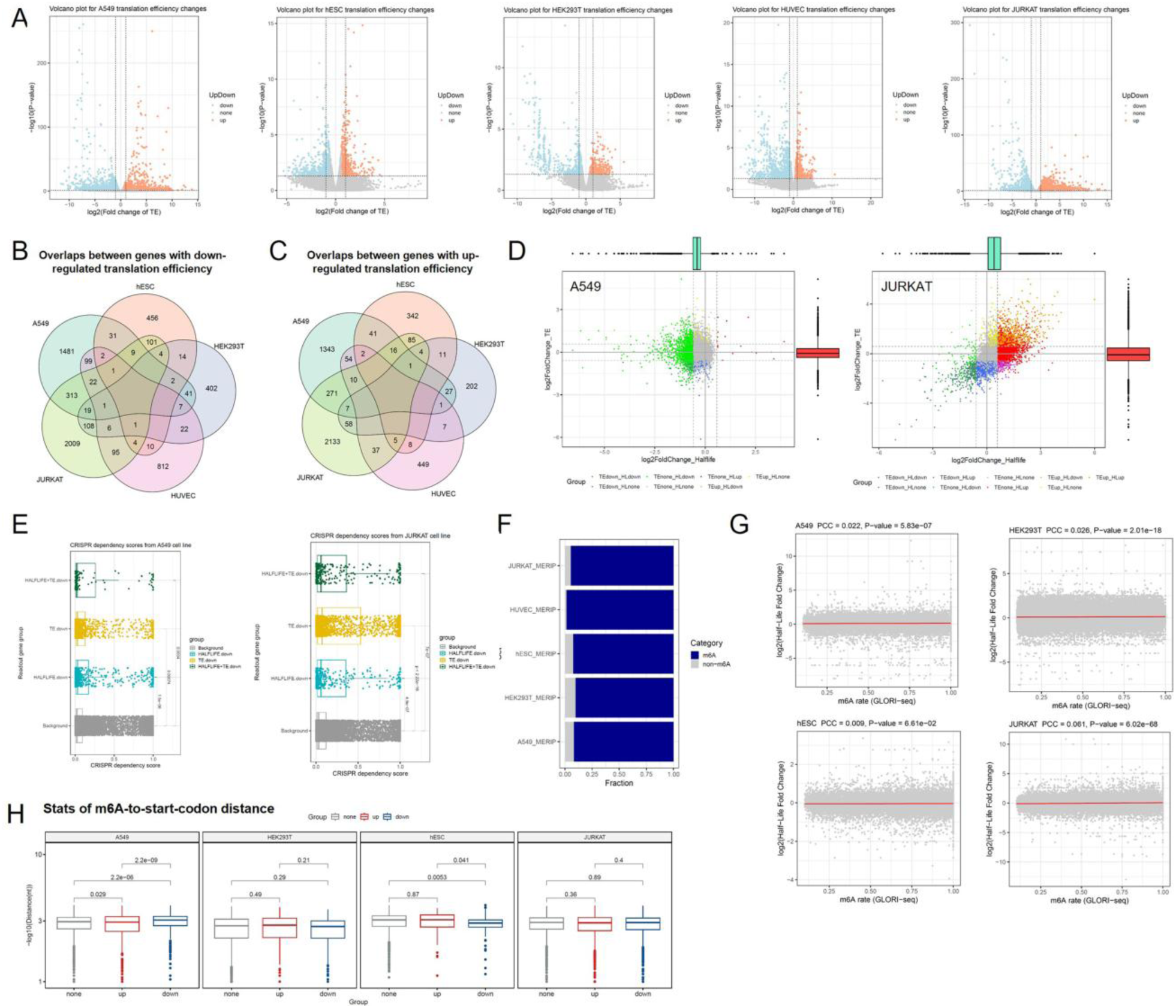
m^6^A methylation regulates translational efficiency in different cell types. (A) Volcano plot of m^6^A-mediated changes in translational efficiency. The comparison was performed between m^6^A-normal cells (shControl) versus m^6^A-disrupted cells (shMETTL3) as the background. (B-C) Venn diagram of translational efficiency down-regulated (B) and up-regulated (C) genes in different cell types. (D) Scatter plot showing translational efficiency versus half-life changes in A549 and JURKAT cells. (E) CRISPR-based gene importance scores of genes with half-life and/or translation efficiency changes in cancer cell lines A549 and JURKAT compared to genome background. (F) Percentage of translation efficiency change genes with m^6^A methylation peaks identified by MeRIP-seq assay on the corresponding cell types. (G) Correlation between fold change in translation efficiency (in absolute values) and m^6^A methylation levels assessed by GLORI. (H) Box plots comparing the minimum m^6^A-site-to-strart-codon distance between genes showing different translational efficiency changes.

The ultra-deep sequencing enabled extensive detection of alternative splicing events, where the m^6^A-mediated alternative splicing was mainly manifested as exon skipping, at both the event and the gene level (**Figure 3A**). Therefore, subsequent analyses of alternative splicing were focused on exon skipping. Generally, an increase of exon inclusion ratio (Psi) could be observed in m^6^A-normal cells (**Figure 3B**), consistent with previous studies ^13,43^. Genes showed altered exon inclusion were less shared between different cell types (**Figure 3C-D**). Basic functions like organelle organization and DNA repair are enriched for these genes (**Supplementary Figure S4**). However, it is noteworthy that only a small proportion of these exons had known m^6^A methylation sites in their vicinity, suggesting that there might be a greater influence of indirect factors on the observed exon inclusion changes (**Figure 3E and Supplementary Figure S2H**). To reduce the influence of indirect factors, for the subsequent analysis we only considered exons with m^6^A methylation sites within 10kb flanking the exon boundary. Among these exons, the m^6^A-to-exon-boundary distances were only weakly or not significantly correlated with the changes in exon inclusion ratios (**Figure 3F**). In addition, GLORI-estimated methylation levels were only weakly correlated or not significantly correlated with changes in exon inclusion ratio (**Figure 3G**), suggesting that simple transcript topological features are not sufficient to predict the alternative splicing readout of m^6^A.

**Figure 3.**
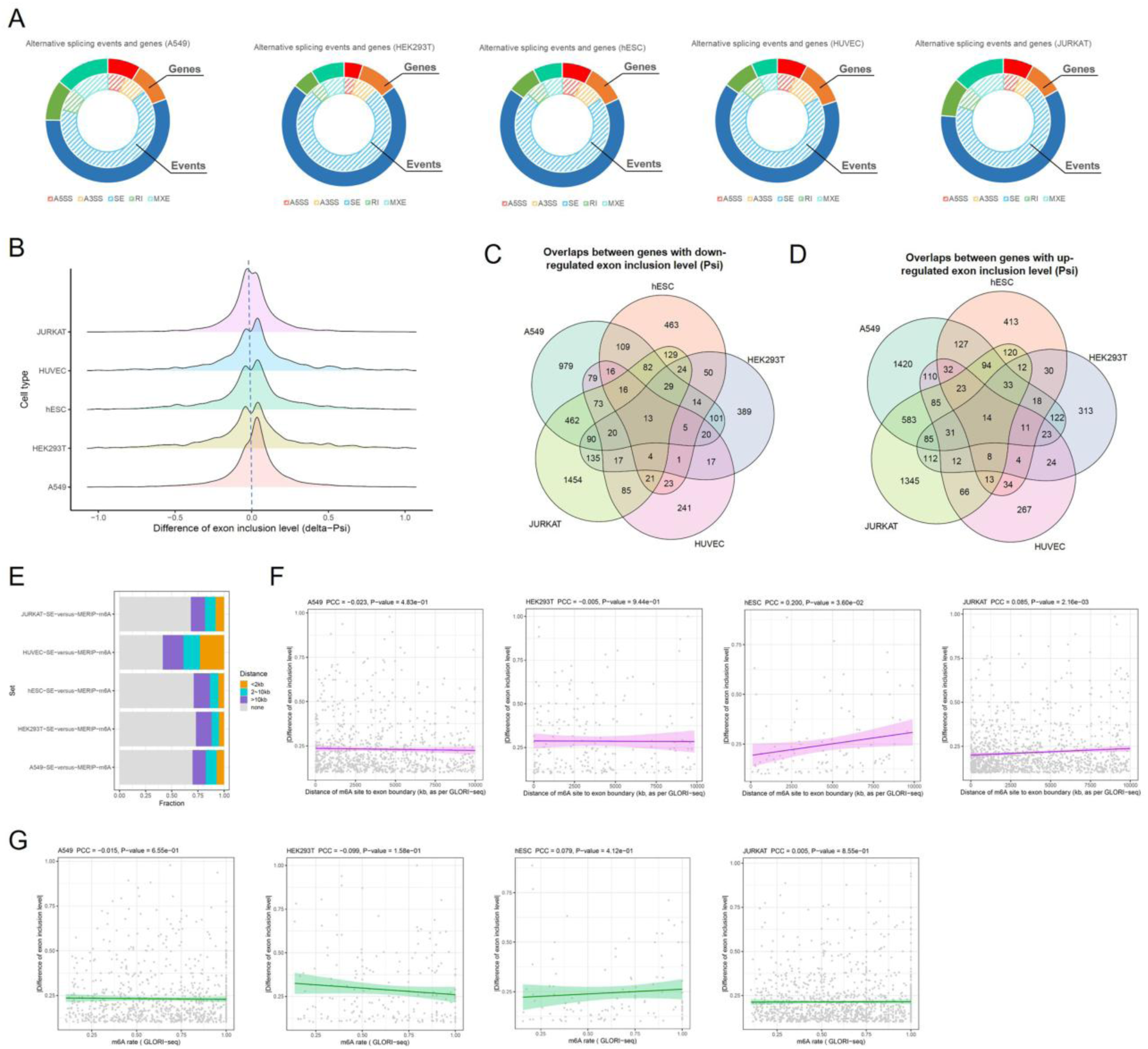
m^6^A methylation regulates alternative splicing in different cell types. (A) Circular plot summarizing different types of alternative splicing changes mediated by m^6^A. The inner and outer loops representing the percentages of alternative splicing events and genes, respectively. (B) Ridge plot summarizing the m^6^A-mediated exon inclusion ratio (Psi) changes. The comparison was performed between m^6^A-normal cells (shControl) versus m^6^A-disrupted cells (shMETTL3) as the background. (C-D) Venn diagram of exon inclusion ratio down-regulated (C) and up-regulated (D) genes in different cell types. (E) Percentage of exons showing exon inclusion ratio changes with m^6^A methylation peaks (identified by MeRIP-seq assay on the corresponding cell types) near their exon boundaries. (F) Correlation between changes in exon inclusion ratio (in absolute values) and m^6^A (within 10 kb to exon boundary) methylation levels assessed by GLORI. (G) Correlation between changes in exon inclusion ratio (in absolute values) and the distances from m^6^A methylation sites to exon boundaries.

### Predicting m^6^A readouts based on the RBP binding context

Given that m^6^A methylation level and topology could not accurately predict m^6^A functional readout, we paid more attention on the genomic context, especially the RBP binding context of the m^6^A sites. To this end, we systematically collected and re-analyzed high-throughput RBP binding site data from public databases (**Supplementary Table S5**) to comprehensively evaluate the associations between m^6^A sites and RBP binding sites. To ensure applicability, we used MeRIP-seq-based m^6^A site annotation here (see Methods; **Supplementary Table S4**). The score of the closest RBP binding site and the genomic/transcript distance were calculated as the RBP features of an m^6^A site. We also introduced the distribution features of m^6^A sites in transcripts, tissue expression level, gene importance score and k-mer features to better characterize m^6^A sites and their target genes. The above features were inputted into the XGBoost model to build the m^6^A readout predictor, hereafter referred as Reading-m6A. Reading-m6A could deal with five readout categories: half-life down-regulation, translational efficiency down-regulation and up-regulation, and exon inclusion ratio decrease or increase. The half-life up-regulation cases were not considered due to the limited number of this type of genes. Because of the obvious differences in m^6^A readout between different cells, each cell type was modeled separately and the prediction accuracy of the model was assessed using ten-fold cross-validation. The results showed that the overall prediction performance of the model was acceptable, with an average AUROC of 0.8, especially for half-life and alternative splicing readouts (**Figure 4A-E**). To further test this modeling approach, we also introduced translational efficiency readout data derived from public Ribo-seq datasets. Four cell lines (HeLa, HepG2, Huh7 and MOLM13) were included. and the predictive performance of the model is equally acceptable in these cells except MOLM13 (**Fig. 4B, C**). Compared to other classical machine learning frameworks, XGBoost showed the best performance for most cases (**Supplementary Figure S5A**); and its performance is insensitive to hyperparameter optimization, with at most 0.01 AUC gain after hyperparameter optimization (**Supplementary Figure S5B**). We also checked whether considering GLORI-based m^6^A sites, especially their m^6^A modification rates, could result in better prediction performance. The results demonstrated that replacing MeRIP-seq-based m^6^A site annotation with GLORI-based ones could benefit the performance in some cases but reduce the performance in other cases, even the highly comprehensive GLORI profile from HEK293T was adopted, while including m^6^A modification rate features show marginal influence to the performance (**Supplementary Figure S5C**). Therefore, we kept MeRIP-seq-based m^6^A site annotation as the primary source to build our models.

**Figure 4.**
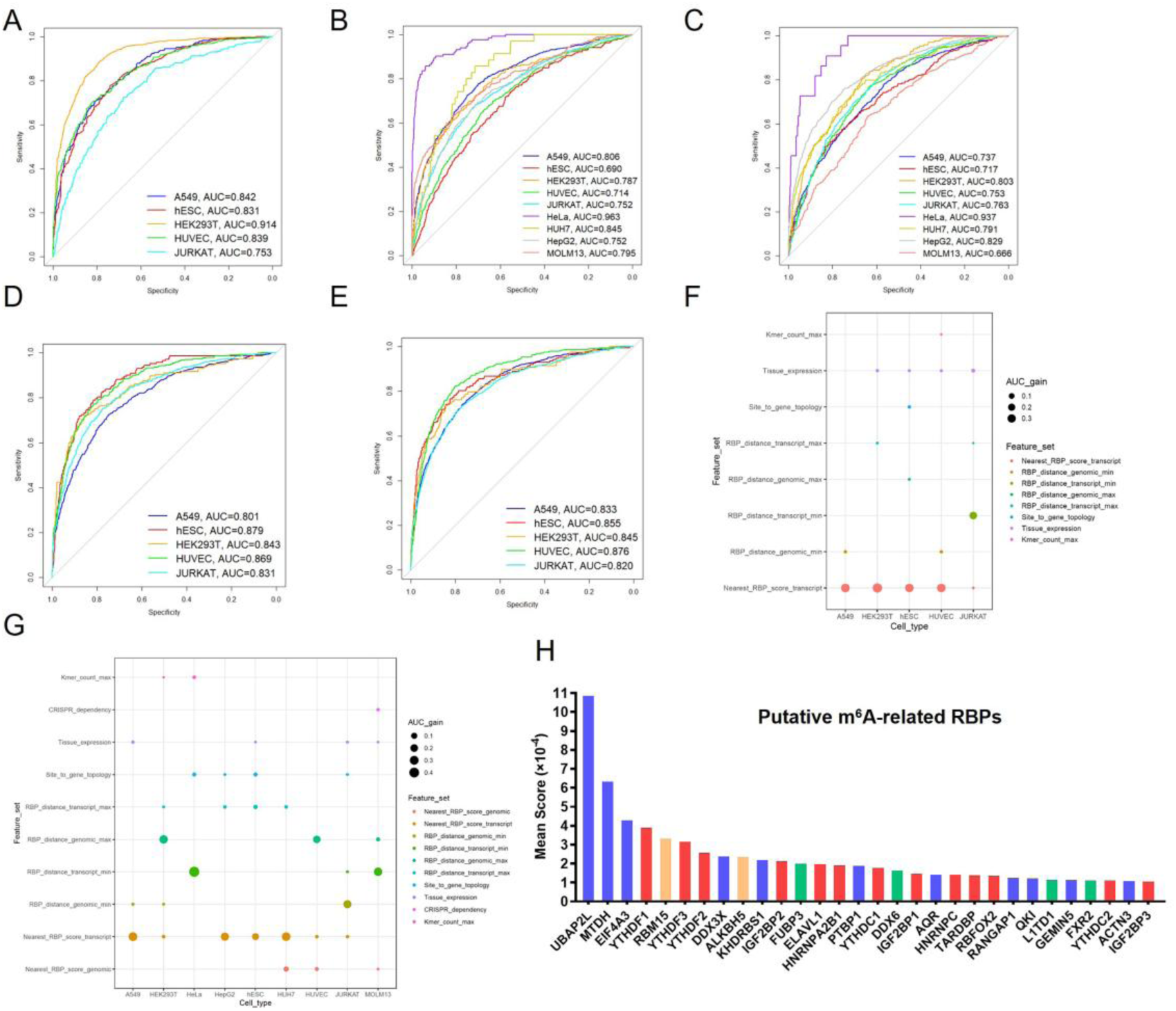
Machine learning models of m^6^A readouts and their top predictive features. (A-E) ROC curve assessing the performances of machine learning models (termed as Reading-m^6^A) for predicting the m^6^A readout effects in terms of half-life down-regulated genes (A), translational efficiency down-regulated genes (B), translational efficiency up-regulated genes (C), exon inclusion ratio down-regulated exons (D), and exon inclusion ratio up-regulated exons (E) in different cell types. (F) Bubble plot summarizing the performance contributions (as evaluated by the gain of AUC) of different feature sets for predicting half-life down-regulated genes. (G) Bubble plot summarizing the performance contributions of different feature sets for predicting translational efficiency down-regulated genes. (H) Bar plot summarizing the RNA binding proteins (RBPs) with high feature importance score in the machine learning models, where red color indicates known m^6^A readers, orange color indicates known m^6^A writers/erasers, green color indicates the newly identified m^6^A reader in this study (FUBP3, DDX6, L1TD1 and FXR2), and blue color indicates the RBPs showing no significant m^6^A binding ability in our RNA pull-down assay.

One of the main reasons we used the XGBoost framework for modeling was its good interpretability. We found that the RBP-based feature sets indeed ranked the best for their contribution to AUC. For half-life and translational efficiency readout predictions, the binding scores of RBP-binding sites near the m^6^A site were the most important feature set (**Figure 4F-G** and **Supplementary Figure S5D**). While the transcript of genomic distance between m^6^A and RBP-binding site ranked top in exon inclusion readout predictions (**Supplementary Figure S5E-F**). Gene topology features and tissue expression contributed to the performance to a much smaller extent. K-mer features showed only marginal or no contribution to AUC, implying the context features related to m^6^A readouts were largely covered by current RBP binding site data. Moreover, further analysis of the importance scores of individual RBP features revealed that some known m^6^A readers had high importance scores, while other important RBP features suggested putative novel m^6^A readers, for which we will be experimentally validated in the next sections (**Figure 4H**).

### M^6^A readout-related RBPs as the novel m^6^A readers

We noticed some RBPs with high feature importance in our m^6^A readout models have been reported as m^6^A readers ^4^. These RBPs included typical m^6^A readers such as YTHDF1/2/3, YTHDC1/2 and IGF2BPs as well as some recently discovered m^6^A readers like TARDBP and ELAVL1 (**Figure 4H**). To further screened novel m^6^A-binding proteins, we used methylated single-stranded RNA bait (ss-m^6^A, with the consensus sequence GG(m^6^A)CU) or unmethylated control RNA (ss-A) to perform RNA pull-down assay, where several known m^6^A readers (YTHDF1, YTHDC2, RBFOX2, TARDBP) were included as the positive controls (**Figure 5A-B** and **Supplementary Figure S6**). The pull-down screening was mainly performed in A549 cells, and the results showed that DDX6, FUBP3 and FXR2 could strongly bind to the methylated bait instead of the unmethylated control (**Figure 5A**), and their binding to m^6^A was also confirmed in hESCs (**Figure 5B**). Because L1TD1 has no expression in A549 cells, we could only validate L1TD1 binding to methylated bait in hESCs only (**Figure 5B**).

**Figure 5.**
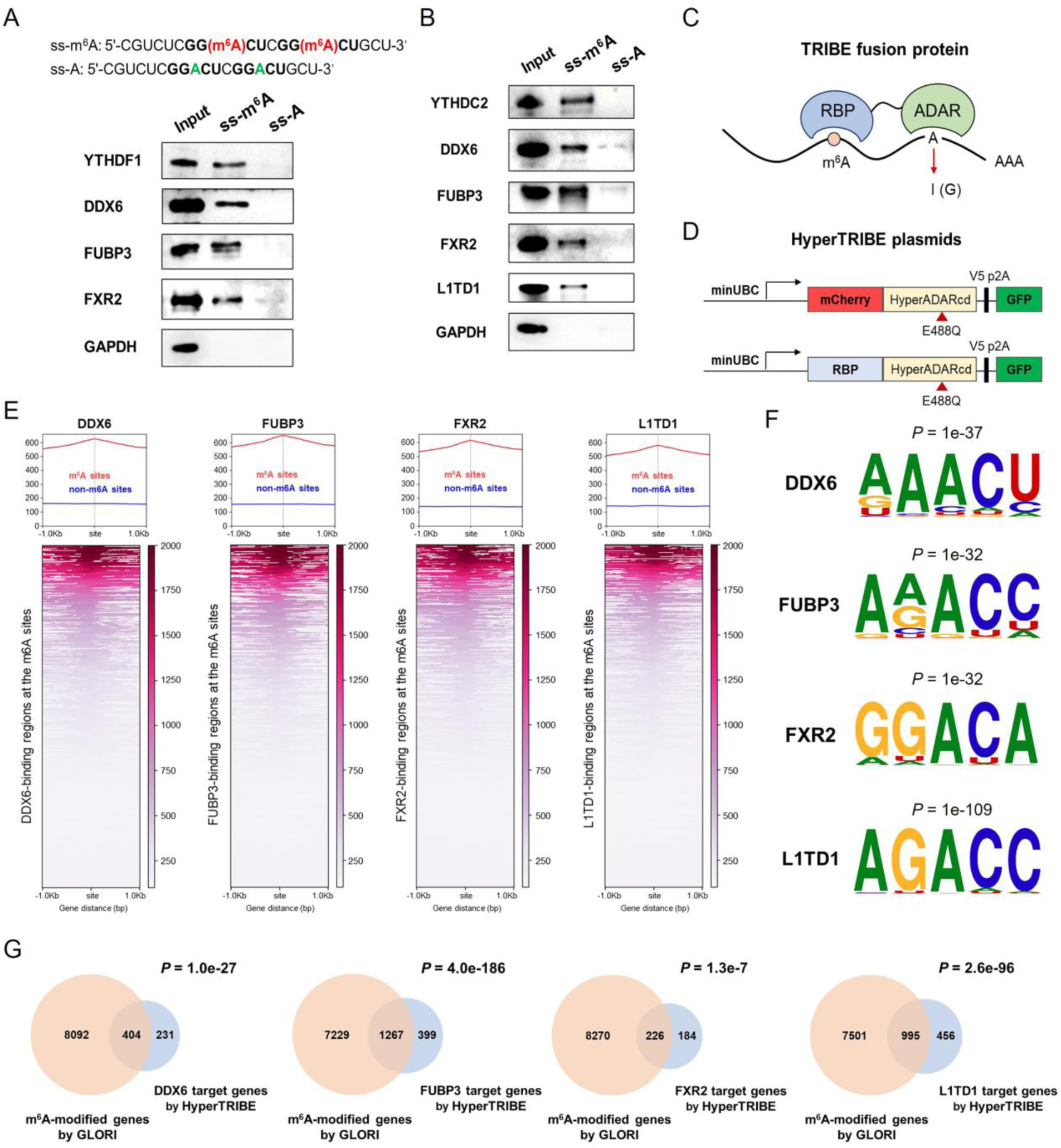
Identification of m^6^A readers in hESCs. (A) RNA pull-down assays that verify the binding of RBPs to m^6^A in A549 cells. (B) RNA pull-down assays that verify the binding of RBPs to m^6^A in hESCs. (C) Overview of principle of the TRIBE/HyperTRIBE for transcriptome-wide identification of RBP binding sites. (D) Diagram of plasmid constructs for HyperTRIBE. (E) Heatmap showing the distribution of HyperTRIBE-identified RBP binding sites near the m^6^A methylation sites determined by GLORI in hESCs. (F) DRACH-like motifs in the proximal sequences of RBP binding sites in hESCs. (G) Venn diagram showing the intersection between RBP target genes identified by HyperTRIBE and m^6^A target genes identified by GLORI in hESCs.

We further investigated the transcriptome-wide binding preference of these candidate readers toward m^6^A sites by using TRIBE/HyperTRIBE technique ^37,44,45^. TRIBE involves fusing the target RBP to the catalytic domain of the RNA editing enzyme ADAR. When the fusion proteins are expressed in target cells, the A-to-I editing at residues in proximity to RBP binding sites was catalyzed by ADAR, providing a signal detectable in conventional RNA-seq (**Figure 5C**). HyperTRIBE exploits a mutated version of ADAR domain to improve its activity thus enhance the A-to-I signals ^37,45^. To demonstrate the feasibility of HyperTRIBE approach, we firstly applied this technique to well-known m^6^A reader YTHDF1 in HEK293T cells (**Figure 5D**). After applying stringent thresholds and subtracting negative controls (see Methods), over 100,000 sites targeted by YTHDF1-ADAR fusion protein were observed. *De novo* motif analysis revealed that YTHDF1 significantly bind to the GGACU consensus sequence, which is a sub-motif of the DRACH m^6^A consensus motif (**Supplementary Figure S7A**). Over 90% YTHDF1 target genes significantly overlapped with known m^6^A-modified genes identified by GLORI (P < 1.0e-200, Fisher’s exact test; **Supplementary Figure S7B**). Moreover, the majority of YTHDF1 target genes overlapped with those identified by the previous TRIBE assay ^9^ (**Supplementary Figure S7C**), and with known m^6^A target genes from MeRIP-seq assay ^46^ (**Supplementary Figure S7D**). These results together demonstrate the feasibility of HyperTRIBE.

Then we applied HyperTRIBE to DDX6, FUBP3, FXR2 and L1TD1 proteins in hESCs. The enrichment heatmaps and profiles showed that DDX6, FUBP3, FXR2 and L1TD1-binding sites were all notably enriched near m^6^A sites rather than non-m^6^A sites (**Figure 5E**). The *de novo* motif analysis revealed that DDX6, FUBP3, FXR2 and L1TD1-binding regions significantly enriched for the DRACH-like motifs (**Figure 5F**). When comparing DDX6, FUBP3, FXR2 and L1TD1 target genes with the m^6^A target genes (identified either by our GLORI profiling or by MeRIP-seq), significant overlaps could be observed (**Figure 5G** and **Supplementary Figure S7E-H**). Taken together, these results indicated that DDX6, FUBP3, FXR2 and L1TD1 are novel m^6^A readers in hESCs.

### Effect of m^6^A readers on early differentiation in hESCs

The regulatory function of DDX6 in hESCs has been clarified ^47^, but the functions of FUBP3, FXR2 and L1TD1 in hESCs have not been fully elucidated. To investigate potential roles for FUBP3, FXR2 and L1TD1 in self-renewal and early differentiation of hESCs, we constructed *FUBP3*, *FXR2* and *L1TD1* knockout (KO) hESCs by using the CRISPR/Cas9 gene-editing approach. The validty of KO constructs were confirmed by Western blot and Sanger sequencing (**Supplementary Figure S8**). We first evaluated the effects of these m^6^A readers on hESC self-renewal, but no significant differences in cell morphology between *FUBP3*, *FXR2* and *L1TD1* KO hESCs and wild-type hESCs were observed (**Figure 6A**). Besides, compared to wild-type hESCs, the protein expressions of core pluripotent markers (OCT4, SOX2 and NANOG) did not have significant changes in *FUBP3*, *FXR2* and *L1TD1* KO cells (**Supplementary Figure S9A-B**). RT-qPCR analysis also found no significantly differences between FUBP3, FXR2 and L1TD1 KO hESCs and wild-type hESCs in the expression of core pluripotent markers (**Supplementary Figure S9C**). These results revealed that FUBP3, FXR2 and L1TD1 had no significant impact on the expression of core pluripotent genes in hESCs. We also employed the CCK8 and ALP staining assays to investigate the effects of FUBP3, FXR2 and L1TD1 on stem cell proliferation. CCK8 assay showed that the growth rate of *L1TD1* KO and *FXR2* KO cells was changed at 24-48 h; but when extending to 72-120 h after cell inoculation, there was no significant difference in proliferation rate between *FUBP3*, *FXR2* and *L1TD1* KO hESCs and wild-type hESCs (**Figure 6B**). The ALP staining assay also showed no obvious distinction in the number of colonies formed by *FUBP3*, *FXR2* and *L1TD1* KO hESCs in comparison with wild-type hESCs (**Figure 6C** and **Supplementary Figure S9D**), indicating that deletion of *FUBP3*, *FXR2* and *L1TD1* had no significant effect on the proliferation rate of hESCs. Together, these results indicated that m^6^A readers FUBP3, FXR2 and L1TD1 had no prominent impact on self-renewal of hESCs.

**Figure 6.**
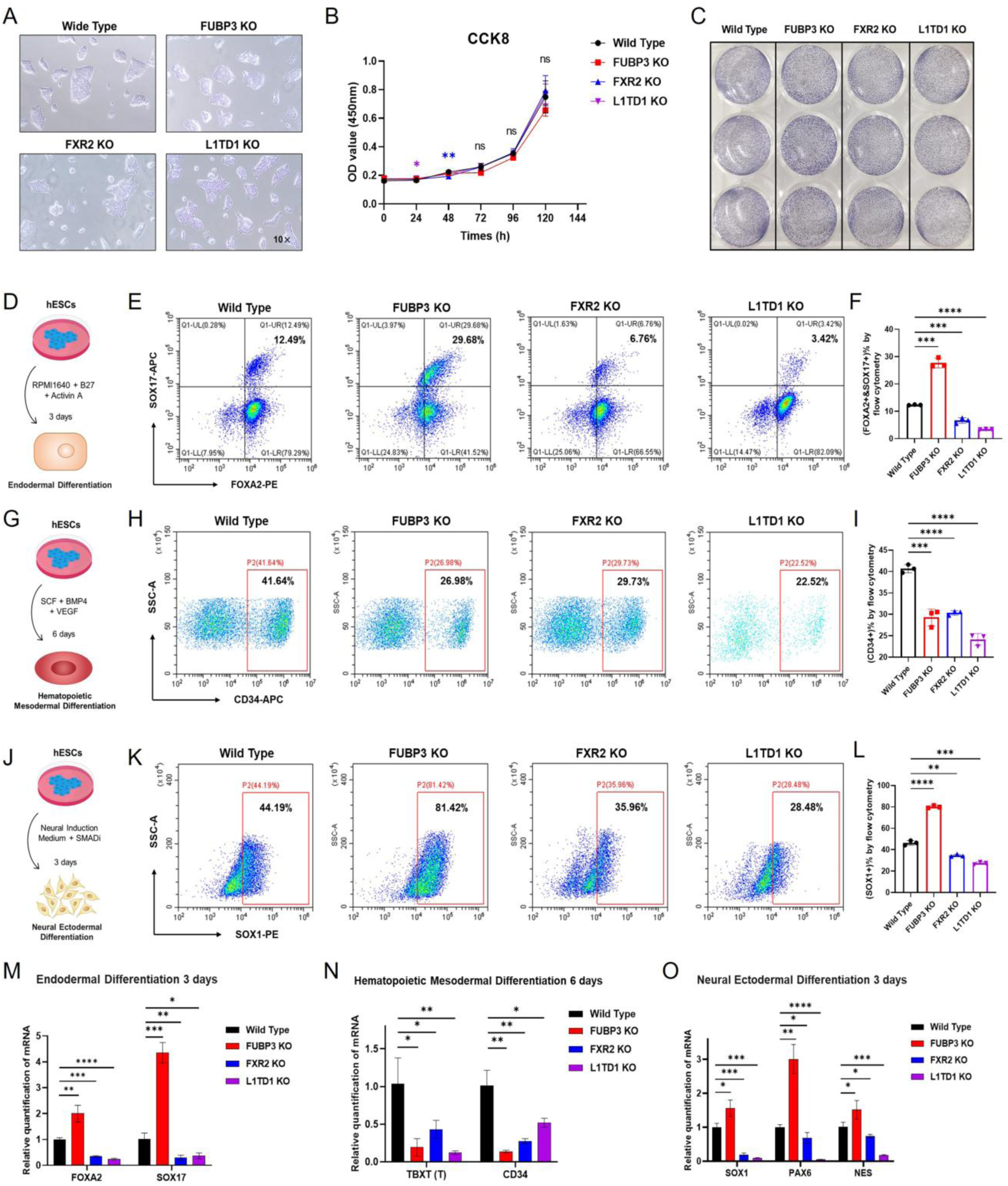
Assessment of m^6^A readers’ regulatory effects on hESC proliferation and differentiation. (A) Representative images of morphology of WT versus *FUBP3*, *FXR2* and *L1TD1* KO hESCs under bright field. (B) Growth curve assessing cell proliferation kinetics of *FUBP3*, *FXR2* and *L1TD1* KO versus WT hESCs. (C) ALP staining assay of WT versus *FUBP3*, *FXR2* and *L1TD1* KO hESCs. (D) Schematic representation of the induction endodermal differentiation experiment of hESCs. (E-F) Flow cytometry and statistical histogram analysis of endodermal differentiation of WT versus *FUBP3*, *FXR2* and *L1TD1* KO hESCs. (G) Schematic representation of the induction hematopoietic mesodermal differentiation experiment of hESCs. (H-I) Flow cytometry and statistical histogram analysis of hematopoietic mesodermal differentiation of WT versus *FUBP3*, *FXR2* and *L1TD1* KO hESCs. (J) Schematic representation of the induction neural ectodermal differentiation experiment of hESCs. (K-L) Flow cytometry and statistical histogram analysis of neural ectodermal differentiation of WT versus *FUBP3*, *FXR2* and *L1TD1* KO hESCs. (M-O) RT-qPCR analysis of definitive endoderm markers SOX17, FOXA2 (M); mesoderm marker TBXT (also known as T) and CD34 (N); and neural-ectoderm markers SOX1, PAX6, NES in WT versus *FUBP3*, *FXR2* and *L1TD1* KO hESCs. (O). Figures in (D), (G) and (J) were drawn by using pictures from Biovisart (https://biovisart.com.cn) and Bioicons (https://bioicons.com/) with permission. For all histograms, * means *P* < 0.05, ** means *P* < 0.01, *** means P < 0.001, **** means P < 0.0001, ns means not significant.

Next, we investigated the effect of m^6^A readers FUBP3, FXR2 and L1TD1 on early differentiation of hESCs by inducing cells into three different germ layers. For endoderm differentiation, Activin A treatment was used in this study. The cells were cultured in RPMI 1640 basic medium supplemented with B27 (**Figure 6D**). After treating with Activin A for 3 days, cells were collected to detect the expression of endodermal markers FOXA2 and SOX17 by flow cytometry and RT-qPCR. The flow cytometry results showed that FOXA2^+^SOX17^+^double positive cells were significantly increased in the *FUBP3* KO cultures compared with wild-type hESCs after endodermal induction for 3 days (**Figure 6E-F**). However, in FXR2 and L1TD1 KO cultures, the proportion of FOXA2^+^SOX17^+^double positive cells was significantly reduced, indicating that loss of *FUBP3* enhances the ability of hESCs to endodermal differentiation, while loss of *FXR2* and *L1TD1* inhibits the ability to differentiate into endoderm (**Figure 6E-F**). As for hematopoietic mesodermal differentiation, cells were seeded on low attachment plates to form EB spheres, and then cultured in STEMdiff APEL^TM^2 medium supplemented with factors SCF, BMP4 and VEGF for 6 days (**Figure 6G**). Flow cytometry results showed that *FUBP3*, *FXR2* and *L1TD1* KO hESCs significantly reduced the proportion of mesodermal marker CD34^+^, indicating that FUBP3, FXR2 and L1TD1 were essential for hematopoietic mesodermal differentiation of hESCs. Deletion of *FUBP3*, *FXR2* and *L1TD1* significantly inhibits the ability to differentiate into mesoderm (**Figure 6H-I**). As for neural ectodermal differentiation, STEMdiff^TM^ neural induction medium with SMADi were used to induce cells to differentiate into neural progenitor cells by monolayer culture protocol (**Figure 6J**). Consistent with the tendency of endoderm differentiation, after neural induction for 3 days, more SOX1^+^positive cells were present in the *FUBP3* KO cultures compared to wild-type group (**Figure 6K-L**). In *FXR2* and *L1TD1* KO cultures, the proportion of SOX1^+^positive cells was significantly declined, indicating that loss of *FUBP3* promotes the ability of hESCs to neural ectodermal differentiation, while loss of *FXR2* and *L1TD1* inhibits the ability to ectodermal differentiation (**Figure 6K-L**).

Furthermore, we detected the mRNA expression level of marker genes of three germ layer through RT-qPCR. Consistent with the flow cytometry results, in endoderm differentiation stage, the mRNA expression levels of FOXA2 and SOX17 were significantly increased in *FUBP3* KO cells but decreased in *FXR2* and *L1TD1* KO cells (**Figure 6M**). During the hematopoietic mesoderm differentiation stage, the mRNA expression levels of TBXT (also known as T) and CD34 were all significantly reduced in *FUBP3*, *FXR2* and *L1TD1* KO cells compared to wild-type group (**Figure 6N**). As for neural ectoderm differentiation stage, the mRNA expression levels of neural-ectoderm markers SOX1, PAX6 and NES (also known as NESTIN) were significantly increased in *FUBP3* KO cells but decreased in *FXR2* and *L1TD1* KO cells (**Figure 6O**). These results indicated that FUBP3 may inhibit the endoderm and ectoderm differentiation, but is important for the mesoderm differentiation. FXR2 and L1TD1 genes are crucial for the differentiation of all three germ layers, and absence may significantly affect the differentiation ability of hESCs. In conclusion, the above results revealed that m^6^A readers play an important role in regulating the early differentiation of hESCs.

## Discussion

In recent years, m^6^A has emerged as a key RNA modification in various biological processes and diseases ^4,48^. Thanks to significant advances in high-throughput sequencing technologies such as MeRIP-seq ^1,2^ and GLORI ^28^, systematic mapping of m^6^A modification sites and their dynamic changes in different tissues, conditions and cell types has become feasible ^3,6–8^. On the other hand, m^6^A modification exhibits diverse post-transcriptional functional readouts, including changes in RNA half-life, in translation efficiency, and in alternative splicing ^49^. Despite the diverse functional readouts of m^6^A, there is a lack of systematic measurement and comparison of m^6^A post-transcriptional functional readouts across multiple cell lines. In fact, an extensive search of the public omics database GEO could only yield a very limited number of cell lines where the m^6^A methylome could be coupled with its readouts. For example, we obtained the m^6^A profiles along with Ribo-seq data for only four cell lines (HepG2, HeLa, Huh7, and MOLM13) with acceptable quality, and these data were introduced as an expanding dataset for translational efficiency readout models. Even for alternative splicing, which can be measured by RNA-seq, few datasets could meet the required sequencing depth for comprehensive alternative splicing identification. To this end, in this study we first selected five representative cell lines (A549, hESC, HEK293T, JURKAT and HUVEC) and performed temporal transcriptome sequencing, Ribo-seq and ultra-deep RNA sequencing on m^6^A-normal (siControl) and m^6^A-disrupted (siMETTL3) cells. Our high-throughput data provide a comprehensive resource for further exploration of the regulatory mechanism as well as the key regulatory targets of m^6^A methylation. Indeed, machine learning modeling of the profiled m^6^A readouts has successfully prioritized novel m^6^A readers from the RBP features of the models, demonstrating the utility of m^6^A readout profiles.

Noticeable cell type specificity of the m^6^A readouts was observed. On the one hand, the m^6^A methylation rate alone is not sufficient to explain this diversity. There are some overlaps of m^6^A target genes between different cell types. However, the overlaps of m^6^A readouts are limited except for some basic pathways. Furthermore, the correlations between GLORI-measured m^6^A modification ratios with the post-transcriptional readouts were compromised, and the inclusion of m^6^A modification ratio did not result in better prediction of m^6^A readouts. Instead, we pay more attention on the association of RBP-binding context of m^6^A sites and m^6^A readouts. Analysis of our readout prediction models suggests that the prediction performance was largely contributed by RBP-based features, either the RBP binding site score near the m^6^A site or the distance between RBP binding sites and m^6^A modification sites. Interestingly, we found that there are few RBPs that significantly contribute to the predictions of readouts, including some well-known m^6^A readers, and screened novel m^6^A readers based on this candidate list. Together, our results emphasize the importance of m^6^A readers, which form a complex regulatory context that controls how the m^6^A modification is interpreted in cells. On the other hand, it is likely that cell type specificity is overestimated here, when compared to the entire repository of human cell types. To be honest, we intentionally selected five cell types from the different cell clades that show distinctive gene expression patterns to ensure representativeness of our data at an affordable cost. When focusing on a specific physiological or disease process, our readout analysis pipeline would be applied to similar cell lines to investigate the consensus m^6^A readout targets and mechanisms.

In this study, we screened DDX6, FUBP3, FXR2 and L1TD1 as the novel m^6^A readers and assessed their roles in regulating pluripotency in hESCs. DDX6 is an RNA helicase belonging to the Dead Box family, which participates in many aspects of RNA metabolism including processing body (P-body) formation ^50^, mRNA storage and decay ^51^, microRNA pathways ^52^, and translational suppression ^51^. DDX6 plays an essential role in regulating self-renewal and differentiation. Di Stefano et al. has reported that DDX6 is required for ESC exit from the pluripotent state by mediating the translational suppression of target mRNAs in P-bodies ^47^. Unlike DDX6, the functions of FUBP3, FXR2 and L1TD1 in ESCs is less explored. FUBP3 is an RNA-binding protein that can bind to the far upstream element (FUSE) of specific RNA molecules, thereby regulating gene expression. In cancer, FUBP3 functions as an oncogene or tumor suppressor gene depending on the type and context of the cancer ^53–55^. However, the role of FUBP3 in cell fate transitions remains largely unclear. Our data in the first time revealed that FUBP3 exerted important impact on early differentiation of three germ lays in hESCs. FXR2 is also an RNA-binding protein related to Fragile X syndrome. FXR2 are highly enriched in mammalian brains, and has important effects on neurogenesis by regulating the proliferation and differentiation of neural progenitors in mice ^56^. In this study, we observed that FXR2 deletion in hESCs impaired early neural development, substantially expanding the scale of FXR2 in neural development regulation. L1TD1 gene was first identified from the expressed sequence tag libraries by Yamanaka group. L1TD1 is abundantly expressed in ESCs and rapidly downregulates upon differentiation ^57^. L1TD1 was reported as a part of the pluripotency interactome network of OCT4, SOX2, and NANOG ^58^. Interestingly, L1TD1 is dispensable for the maintenance of pluripotency in mESCs, despite its specific expression pattern ^57^. We observed that LITD1 deletion had no significant effects on self-renewal in hESCs. Of note, hESCs with LITD1 showed significant defects in early differentiation of three germ layers, meaning that L1TD1 play an essential role in human early embryonic development. Our work uncovered the new roles of FUBP3, FXR2 and L1TD1 in stem cell fate control, and sheds light on post-translational regulation in pluripotency and the importance of RBPs in pluripotent cells.

Our study also has some limitations. Firstly, the scope of cell types included is not sufficient in comparison with the wide spectrum of human cell atlas. Secondly, recent studies have revealed new layers of m^6^A readouts such as the regulatory interplay with chromatin, but this aspect was not covered in current study. Thirdly, the molecular mechanism of the identified m^6^A readers have not been in-depth investigated. Nonetheless, our study has provided useful resource to bridge the gap between m^6^A methylome landscape and its readouts, and prioritizing interesting novel m^6^A readers for stem cell fate control. We expected the future research could expand the range of cell types studied, improve data resources, and delve deeper into the molecular mechanisms underlying m^6^A functional readouts.

## Software availability

Reading-m6A online tool is accessible at http://www.rnanut.net/reading-m6a.

## Data availability

The data can be accessed in Gene Expression Omnibus (https://www.ncbi.nlm.nih.gov/geo) with accessions of GSE179872 (m^6^A readout profiling), GSE290408 (GLORI) and GSE287679, GSE288304 (HyperTRIBE).

## Supplementary Figures

**Supplementary Figure S1.**
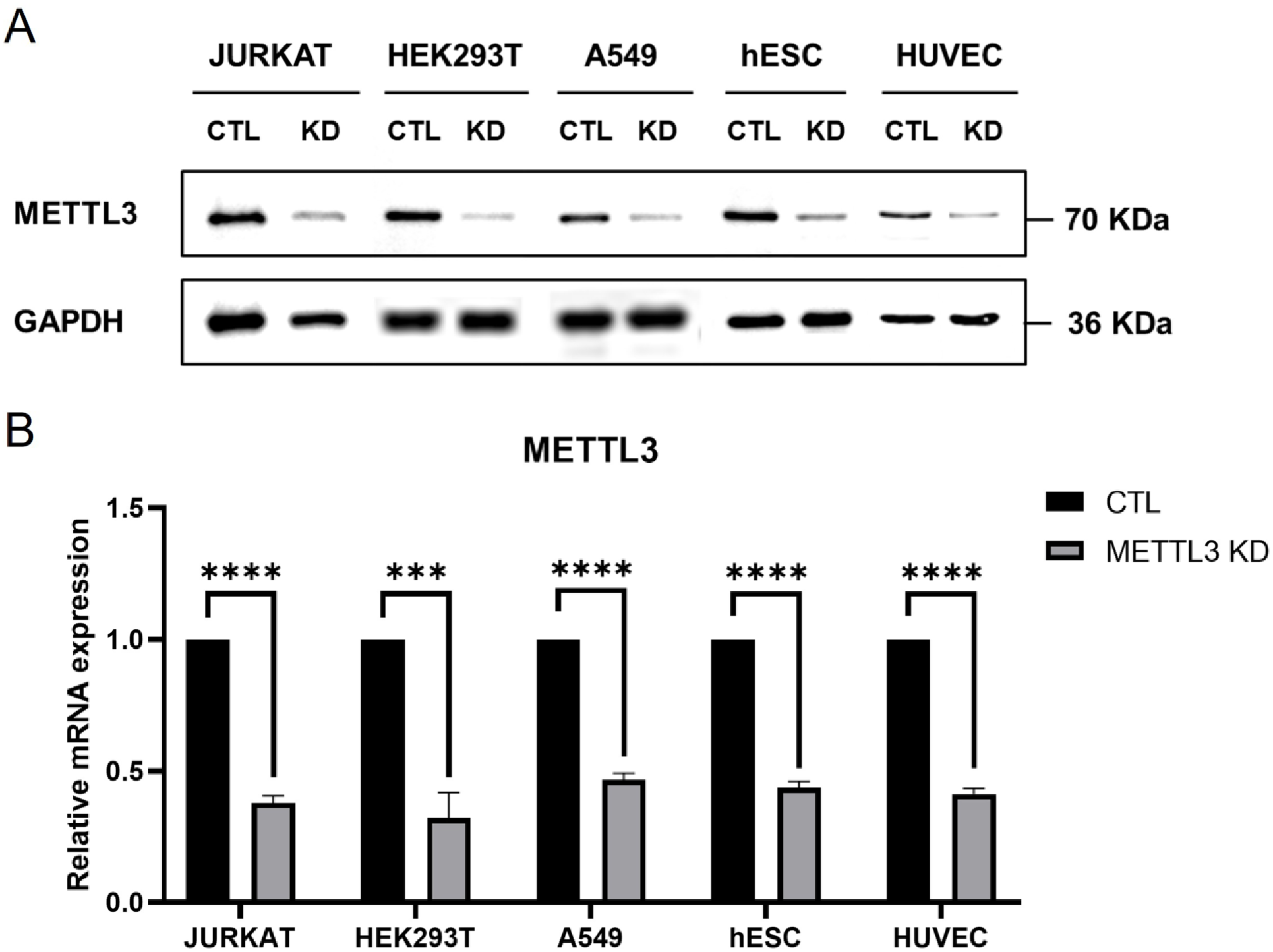
Verification of *METTL3* knockdown effects. (A) Western blot analysis for expression of METTL3 in shControl and METTL3 KD cells. (B) RT-qPCR analysis for the relative expression of *METTL3* mRNA in shControl and METTL3 KD cells. **** means P < 0.0001.

**Supplementary Figure S2.**
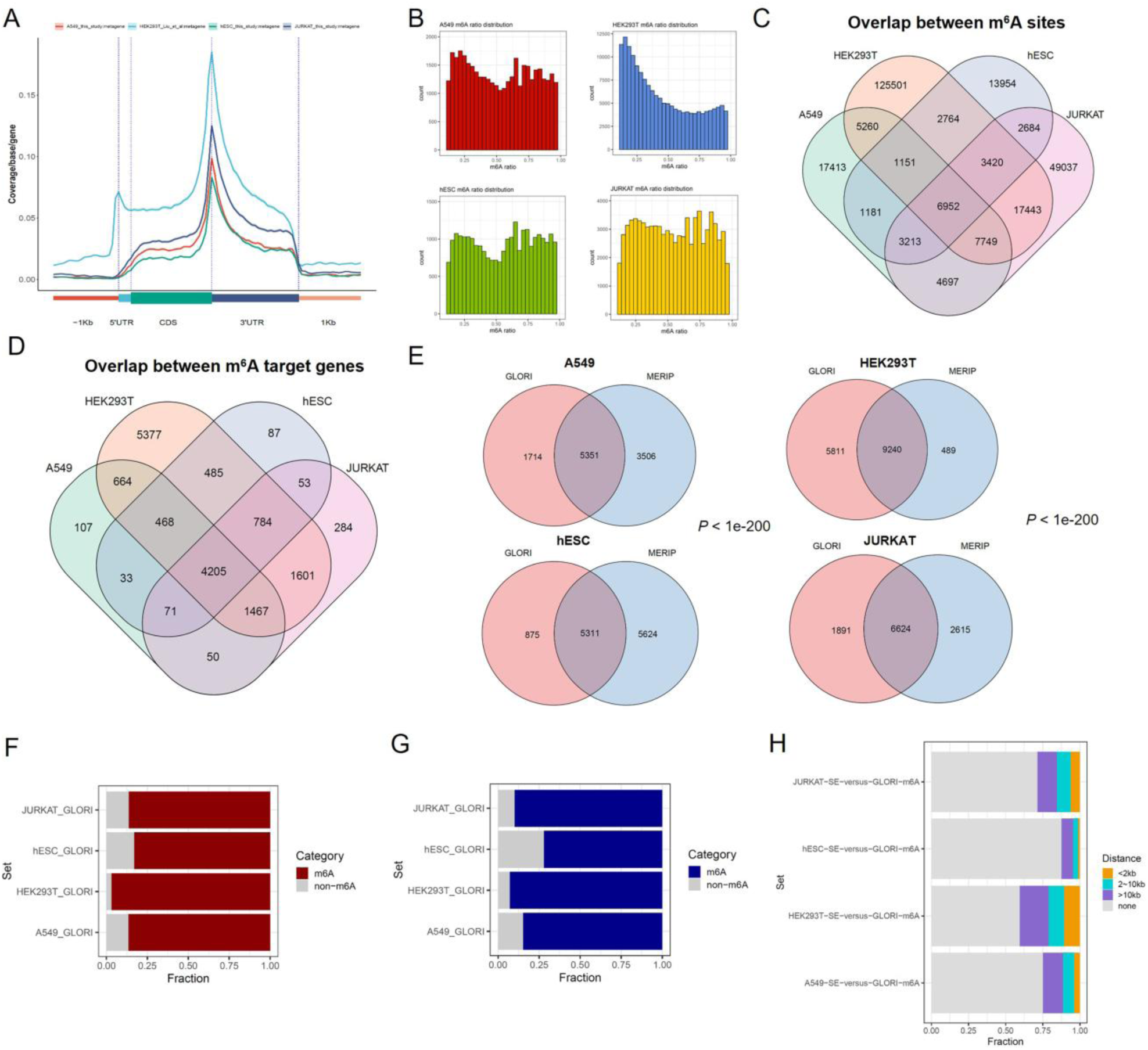
Overview of m^6^A modification sites identified by GLORI. (A) Metagene plot showing the distribution of GLORI-identified m^6^A modification sites in the target genes. Western blot analysis for expression of METTL3 in shControl and shMETTL3 cells. (B) Histogram illustration of m^6^A modification ratio estimated by GLORI. (C) Venn diagram showing the overlap of m^6^A modification sites between different cell types. (D) Venn diagram showing the overlap of m^6^A target genes between different cell types. (E) Venn diagram showing the overlap between m^6^A target genes identified by GLORI and those identified by MeRIP-seq in the corresponding cell types. (F) Percentage of half-life down-regulated genes with m^6^A methylation peaks identified by GLORI on the corresponding cell types. (G) Percentage of genes showing translational efficiency changes with m^6^A methylation peaks identified by GLORI on the corresponding cell types. (H) Percentage of exons showing exon inclusion ratio changes with m^6^A methylation sited identified by GLORI on the corresponding cell types) near their exon boundaries.

**Supplementary Figure S3.**
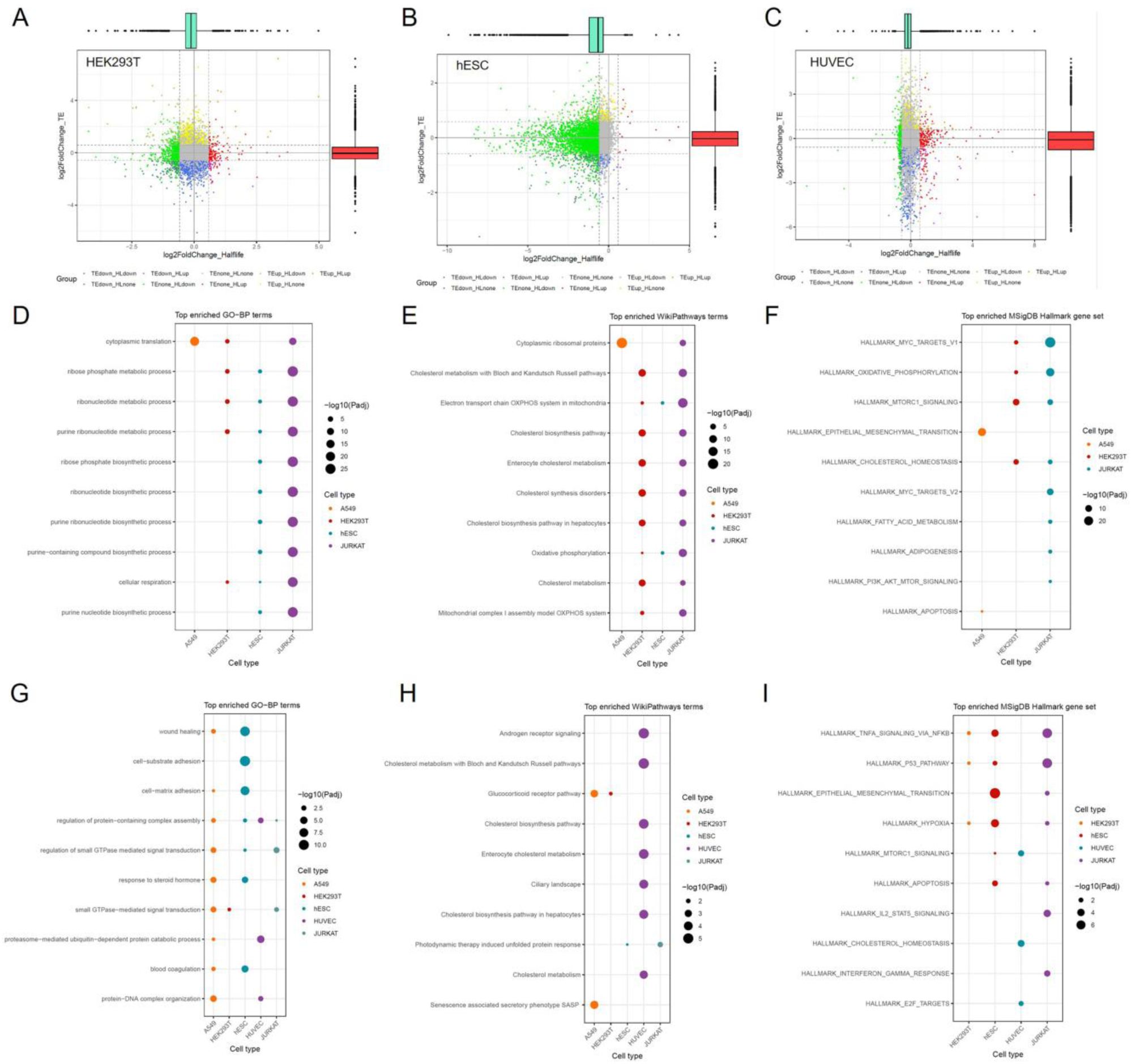
Distribution and functional association of genes showing m^6^A-mediated translational efficiency changes. (A-C) Scatter plot showing translational efficiency versus half-life changes in HEK293T (A), hESC (B) and HUVEC cells (D) Top 10 enriched Gene Ontology-Biological Process (GO-BP) functional terms for translational efficiency down-regulated genes. (E) Top 10 enriched WikiPathways pathways for translational efficiency down-regulated genes. (F) Top 10 enriched HsigDB Hallmark gene set for translational efficiency down-regulated genes. (G-I) The corresponding results for translational efficiency up-regulated genes.

**Supplementary Figure S4.**
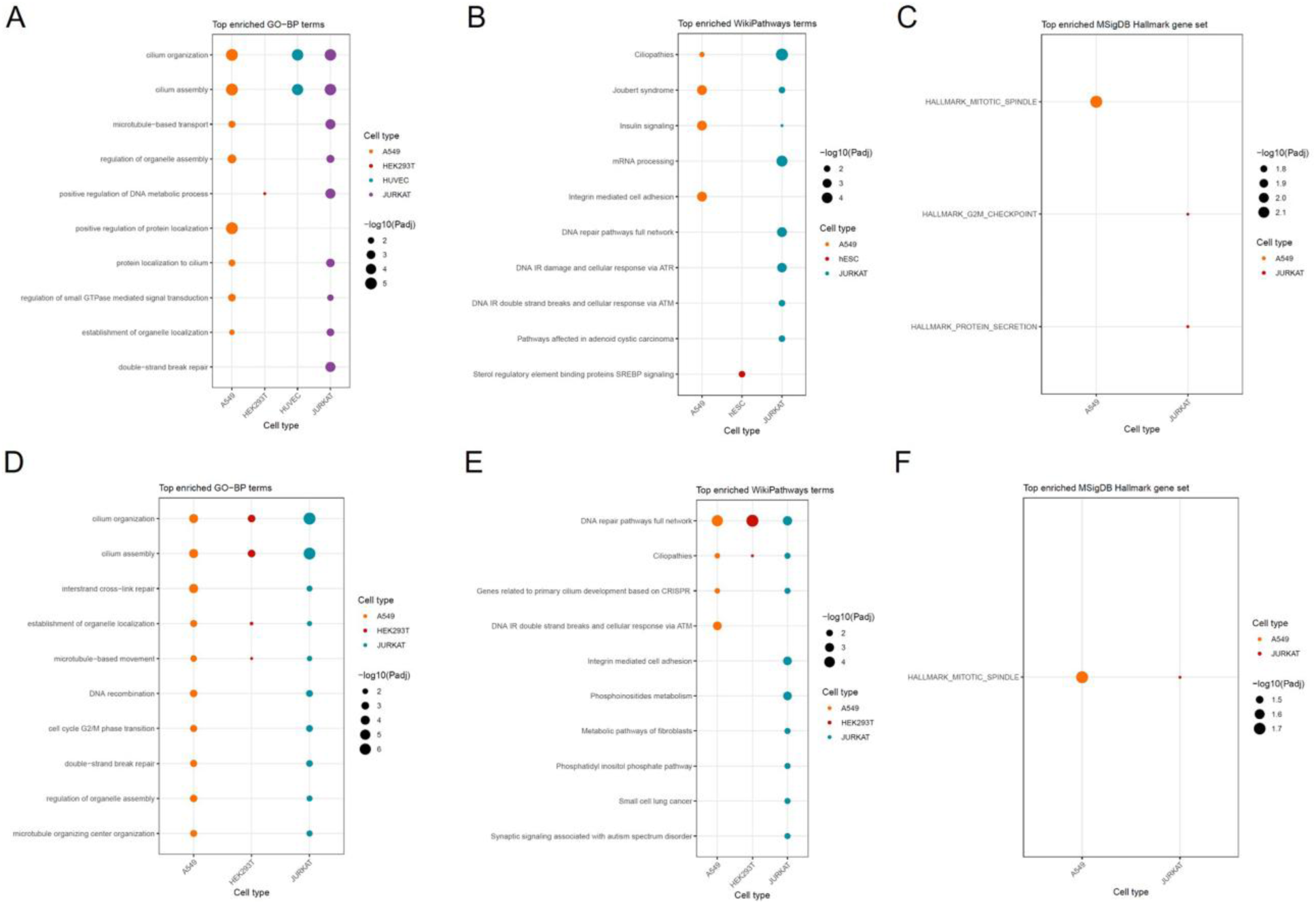
Functional association of genes with exon skipping events. Top 10 enriched GO-BP functional terms for exon inclusion ratio down-regulated genes. (B) Top 10 enriched WikiPathways pathways for exon inclusion ratio down-regulated genes. (C) Top 10 enriched HsigDB Hallmark gene set for exon inclusion ratio down-regulated genes. (D-F) The corresponding results for exon inclusion ratio up-regulated genes.

**Supplementary Figure S5.**
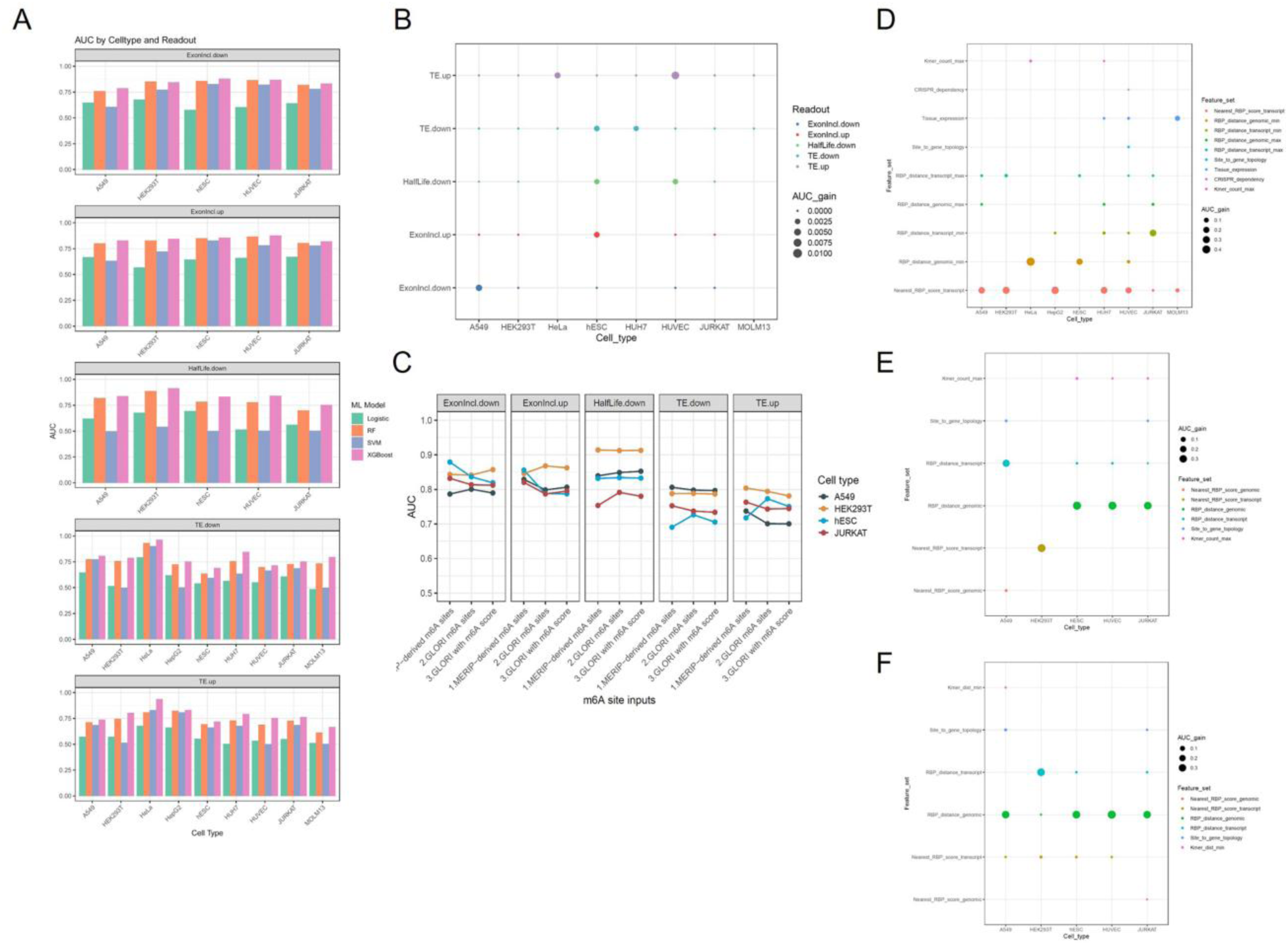
Ablation experiments and feature set contribution of the machine learning predication models of m^6^A readouts. (A) Comparison between XGBoost and other classical machine learning models in predicting different categories of m^6^A readouts. The classical machine learning models here include random forest (RF), logistic regression (Logistic) and support vector machine (SVM). ExonIncl means exon inclusion ratio; TE means translational efficiency. (B) Bubble plot summarizing the gain of AUC after parameter optimization of the XGBoost model. (C) Line plot comparing the performance of XGBoost models with MeRIP-seq-derived m^6^A sites, GLORI-derived m^6^A sites or GLORI-derived m^6^A sites plus the corresponding m^6^A methylation rate. (D) Bubble plot summarizing the performance contributions of different feature sets for predicting translational efficiency up-regulated genes. Bubble plot summarizing the performance contributions of different feature sets for predicting exon inclusion ratio down-regulated exons. (F) Bubble plot summarizing the performance contributions of different feature sets for predicting exon inclusion ratio up-regulated exons.

**Supplementary Figure S6.**
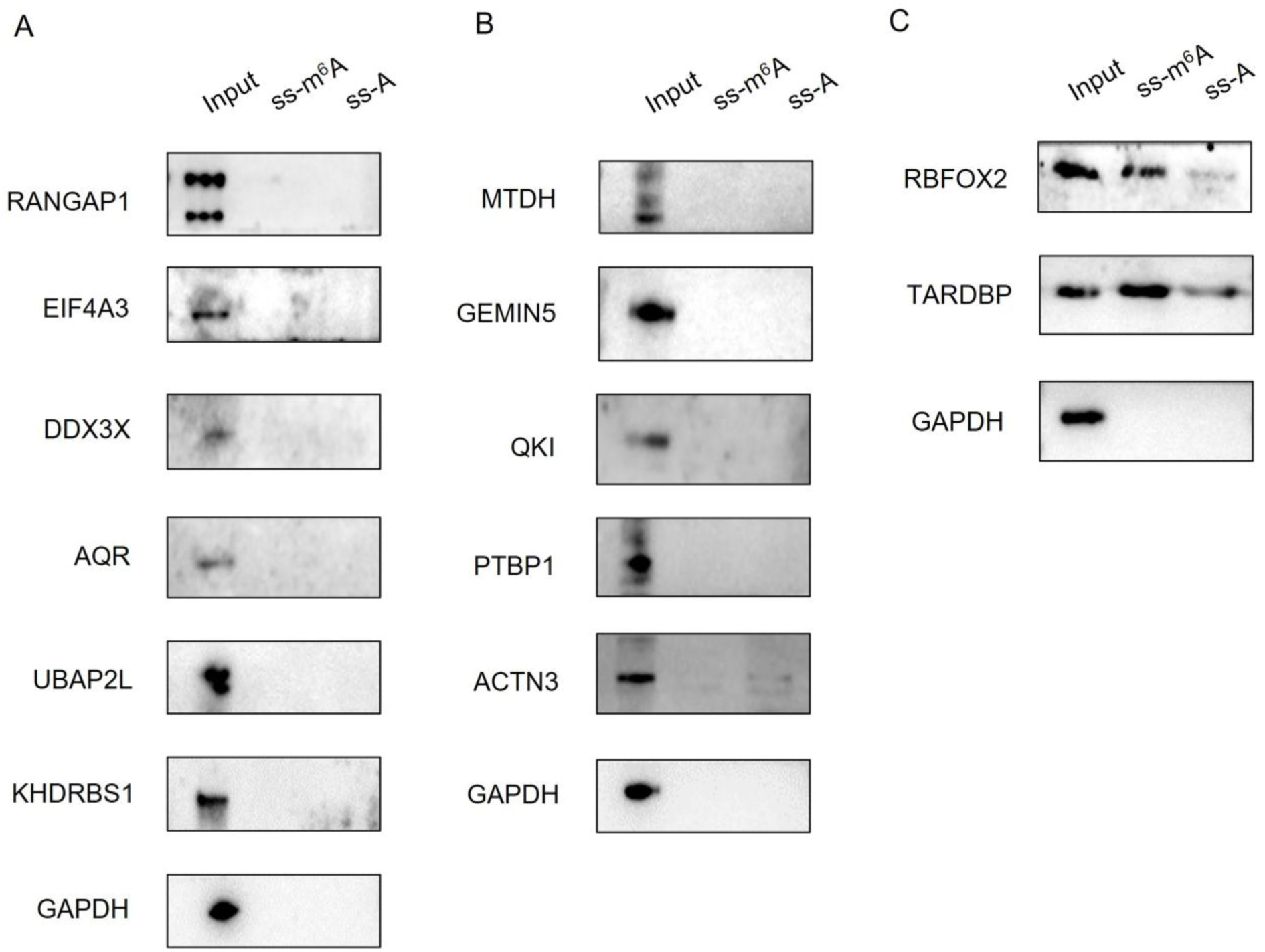
Screening of m^6^A readers by RNA pull-down assay. Three batches of RNA pull-down assay were performed. (A) RNA pull-down assay batch 1. (B) RNA pull-down assay batch 2. (C) RNA pull-down assay batch 3. RBFOX2 and TARDBP (also known as TDP43) are known m^6^A readers that serve as positive controls here.

**Supplementary Figure S7.**
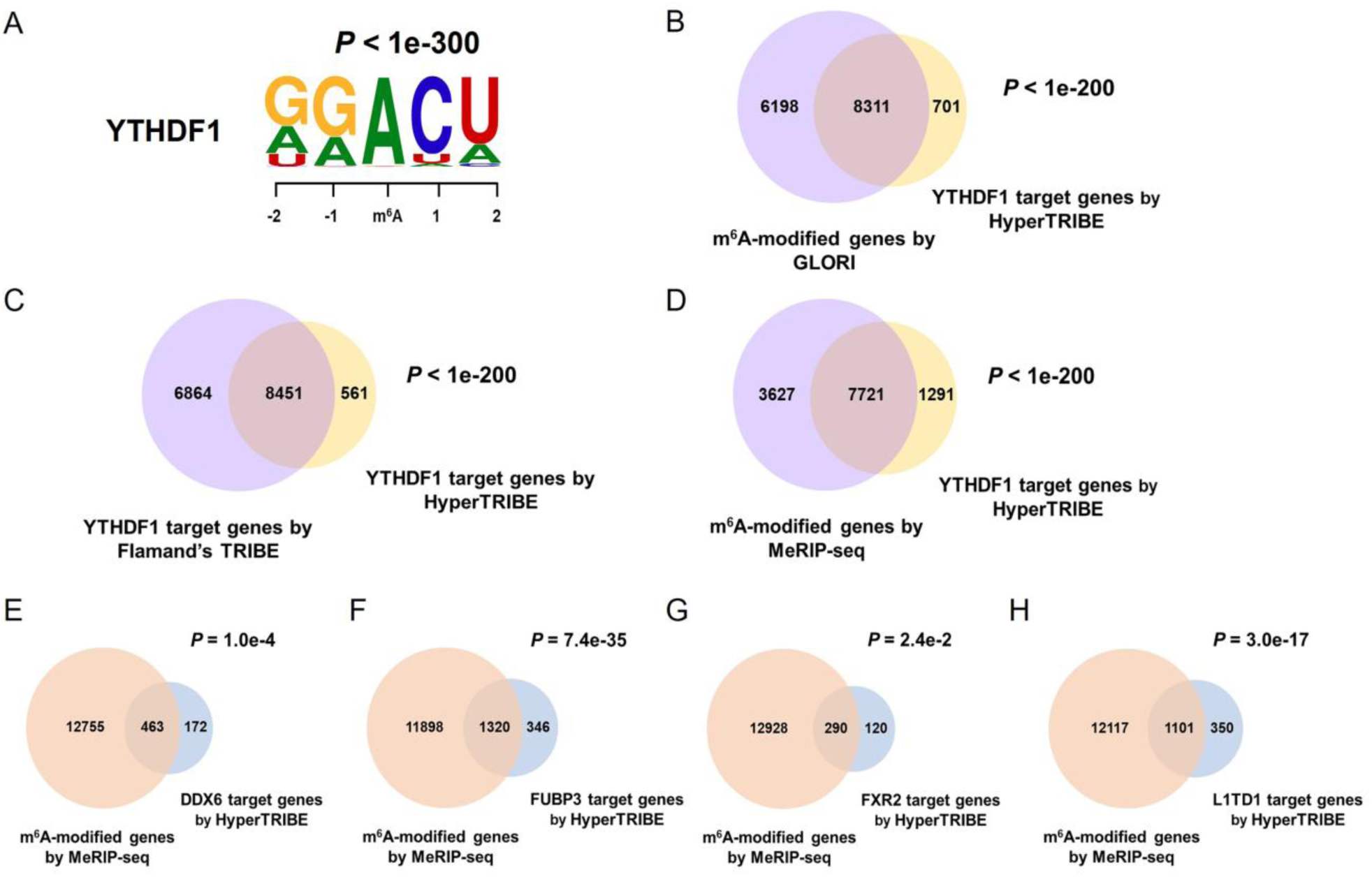
Overlaps between the binding target genes of m^6^A readers and m^6^A modification. HyperTRIBE assay of YTHDF1 in HEK293T was performed as the positive control assay. Then the HyperTRIBE assay was performed in hESCs for the four putative m^6^A readers (DDX6, FUBP3, FXR2 and L1TD1) found in this study. (A) DRACH-like motifs in the proximal sequences of YTHDF1 binding sites in HEK293T. (B) Venn diagram showing the intersection between YTHDF1 target genes identified by HyperTRIBE and m^6^A target genes identified by GLORI in HEK293T. (C) Venn diagram showing the intersection between YTHDF1 target genes identified by HyperTRIBE and those identified by Flamand et al. (D) Venn diagram showing the intersection between YTHDF1 target genes identified by HyperTRIBE and m^6^A target genes identified by MeRIP-seq in HEK293T. (E-H) Venn diagrams showing the intersection between DDX6 (E), FUBP3 (F), FXR2 (G) and L1TD1 (H) target genes identified by HyperTRIBE and m^6^A target genes identified by MeRIP-seq in hESC.

**Supplementary Figure S8.**
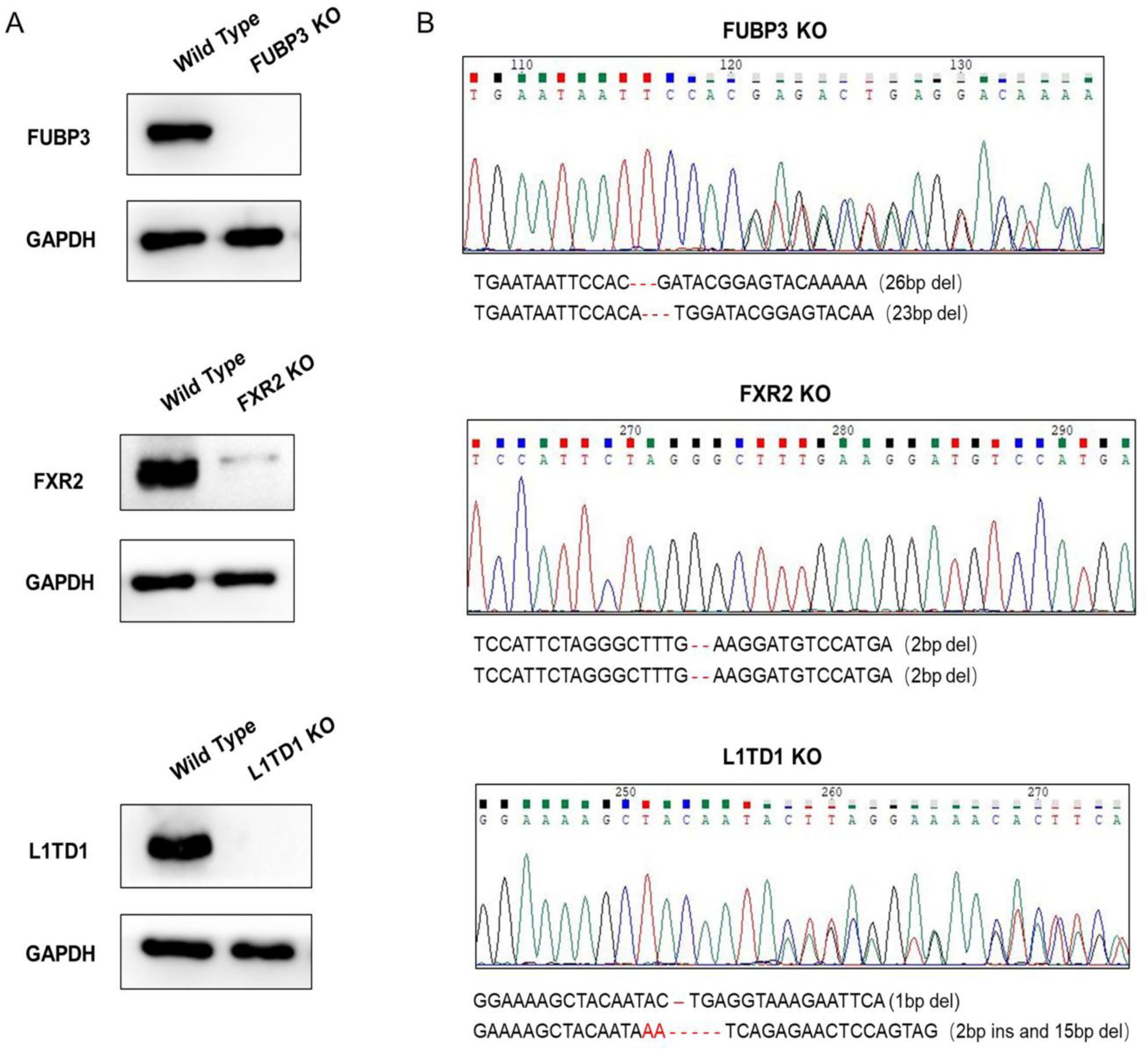
Validation of *FUBP3*, *FXR2* and *L1TD1* KO cell lines by Western blot and sanger sequencing. (A) The validation of *FUBP3*, *FXR2* and *L1TD1* KO cell lines by Western blot. (B) The validation of *FUBP3*, *FXR2* and *L1TD1* KO cell lines by sanger sequencing.

**Supplementary Figure S9.**
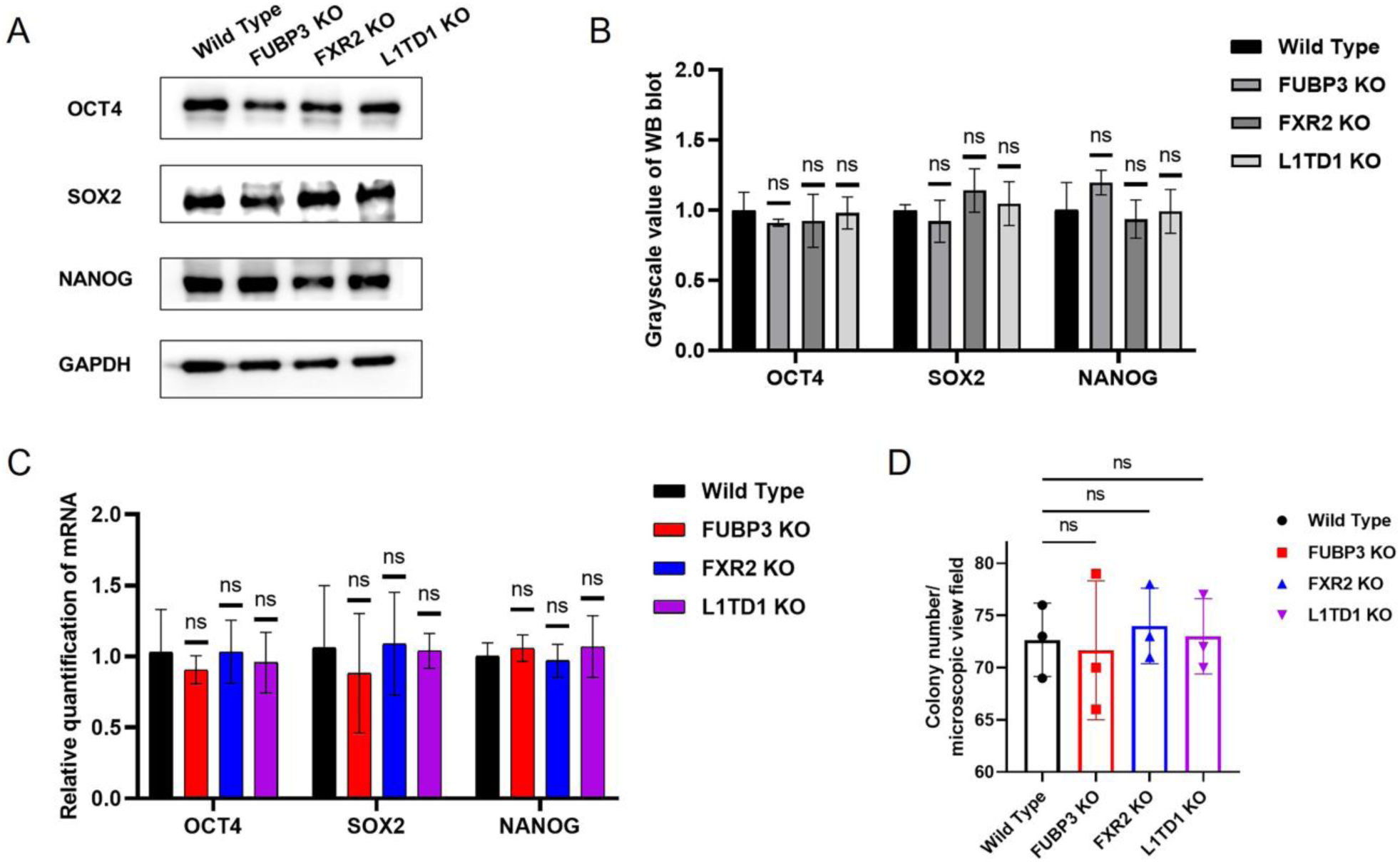
Regulatory effects of m^6^A readers on self-renewal of hESCs. Representative western blot analysis of expression of OCT4, SOX2 and NANOG in WT versus *FUBP3*, *FXR2* and *L1TD1* KO hESCs. (B) The statistics of grayscale value of western blot, ns means not significant. (C) RT-qPCR analysis of OCT4, SOX2, NANOG mRNA expression in WT versus *FUBP3*, *FXR2* and *L1TD1* KO hESCs. ns means not significant. (D) Statistics of the ALP-positive colony numbers from WT versus *FUBP3*, *FXR2* and *L1TD1* KO hESCs. ns means not significant.

